# *Acinetobacter guillouiae* strain isolated from sludge capable of partially degrade polyethylene terephthalate: genomic and biochemical insights

**DOI:** 10.1101/2023.11.05.565377

**Authors:** Naheed Akhtar, Afef Najjari, Cecilia Tullberg, Muhammad Siddique Awan, Zahid Majeed, Carl Grey, Baozhong Zhang, Javier A. Linares-Pastén

## Abstract

The escalating accumulation of plastic waste in terrestrial and aquatic ecosystems profoundly threatens environmental health and biodiversity while impacting human well-being. Recently, many microorganisms capable of degrading polyethylene terephthalate (PET) have been reported, primarily sourced from terrestrial soils and marine environments. Notably, the challenge of PET pollution in aquatic environments has remained a persistent research concern. In this study, we present the isolation and characterization of *Acinetobacter guillouiae* strain I-MWF, obtained from a wastewater treatment plant in Makri, AJK, Pakistan, using molecular phylogenetic analysis based on genome sequencing. Results revealed that this strain exhibits the ability for PET powder degradation, as confirmed by HPLC/LCMS analysis. Furthermore, we conducted whole-genome sequencing using Illumina technology and bioinformatically explored this strain’s potential repertoire of lipase and esterase enzymes. Under optimized conditions of 23°C and pH 7 in mineral salt media with PET powder as the sole organic substrate, *A. guillouiae* I-MWF could degrade partially. Extracellular enzymes yielded PET depolymerization products identified as mono(2- hydroxyethyl) terephthalic acid and terephthalic acid. The sequenced genome of this strain spans 4.61 Mb with a mean G + C content of 38.2%, containing 4,178 coding genes, 71 tRNA, and six rRNA genes. Although no cutinase-like enzymes were identified, our analysis unveiled a diverse array of putative lipases and three esterases, all sharing the typical α/β hydrolase fold. Additionally, comprehensive molecular modelling analysis suggested that some of the 18 identified extracellular hydrolases may be involved in polyester enzymatic depolymerization processes.

## Introduction

Due to the global pollution caused by plastics, mainly single-use plastic products, the research on plastic recycling and degradation has significantly increased in recent years. Biological technologies are emerging as one of the most attractive alternatives to chemical or mechanical processes because of their mild reaction conditions, environmentally friendly procedures, and the potential to become a sustainable solution (Hiraga et al., 2019). The most advantageous aspect of using microorganisms is the possibility of incorporating the products of plastic depolymerisation, that is, monomers, into the central metabolism. Thus, fermentation could turn these monomers into various products of commercial value (Salvador et al., 2019).

Bacteria and fungi produce, so far, the most plastic-active enzymes known, and recently, the first Archaeal PET-degrading enzyme was reported (Perez-Garcia et al., 2023). Bioinformatic studies, in the context of a global microbiome, predicted 30,000 enzymes that could degrade 10 different plastic types (Zrimec et al., 2021). Another bioinformatic study has indicated 504 putative PET active enzymes in bacteria, mainly in the phyla of *Actinobacteria*, *Proteobacteria,* and *Bacteroidetes* (Danso et al., 2018). However, currently, less than 50 enzymes with PET and polyurethane (PUR) hydrolytic activity have been experimentally verified, most being PET-active enzymes (Buchholz et al., 2022).

Plastics are usually recalcitrant to biological activity. The chemical nature, considerable molecular weight and crystallinity are the main obstacles. However, plastics with heteroatom backbones, such as polyesters and polyurethanes, are more accessible to biological attack than those of polyolefinic backbones, such as polyethylene, polypropylene, and others (Wei and Zimmermann, 2017). Polyesters and polyurethanes have ester and amide (urethane) bonds, respectively, widely present in biological molecules. Therefore, organisms commonly produce enzymes to hydrolyse these types of bonds. Although polyethylene terephthalate (PET) is the most studied polyester from the biodegradation and enzymatic depolymerisation perspectives (Taniguchi et al., 2019) there are several challenges. It is known that the high crystallinity of PET blocks the accessibility to the enzymes, and its amorphization is beneficial for enzymatic depolymerisation. Therefore, thermostable PET depolymerising enzymes, active at temperatures close to the glass transition temperature of PET (67 to 81°C), are among the most effective (Marten et al., 2005).

So far, most known enzymes able to hydrolyse PET are serine hydrolases, including cutinases, carboxylesterases and lipases (Lai et al., 2023). PET hydrolase (*Is*PETase) from *Ideonella skaiensis* (Yoshida et al., 2016) and the leaf-branch compost cutinase (LCC) (Tournier et al., 2020) are the most studied PET-active enzymes and, therefore, can be considered model PET hydrolases. These enzymes and related cutinases depolymerise PET into terephthalic acid (TPA), mono(2-hydroxyethyl) terephthalic acid (MHET), bis(2- hydroxyethyl)-TPA (BHET), and ethylene glycol. The product profile can vary depending on the enzyme concentration, reaction time or specific cutinase; however, MHET usually appears as the dominant product, followed by PTA and a minor amount of BHET.

Under this framework, this work aims to explore bacteria from activated aerobic sludge that can use PET as the sole carbon source. For this purpose, a collection of strains was isolated using a two-step approach, consisting of a selection of plastic-colonising microorganisms and microorganisms capable of degrading PET powder in a mineral salt culture medium. One of the isolated organisms, identified as *Acinetobacter guillouiae,* was the most promising strain, partially able to degrade PET powder. This strain, named I-MWF, was isolated from sludge samples collected from a water filtration plant located at Makri, Muzaffarabad, on the Neelum River of Azad Jammu and Kashmir, Pakistan. Household and commercial waste from these urban and rural populations of 4.2 million produce tonnes of plastic waste, which enters riverine water.

In the PlasticDB database (Gambarini et al., 2022) *A. baumannii* strains are registered from landfills, soil, and plastic waste dumping sites involved in PET degradation. However, the reported biodegradation evidence is based on FT-IR, weight loss, SEM, or CO_2_ production. In this work, we report the enzymatic activity of PET from extracellular extracts obtained during the cultivation of *A. guillouiae* I-MWF in PET powder. HPLC-UV/MS confirmed the PET enzymatic depolymerisation activity and products, while FT-IR monitored microbial activity on PET. In addition, the *A. guillouiae* I-MWF genome was sequenced, and the putative enzymes involved in polyester degradation were predicted in silico by modelling the 3D structures.

## Materials and methods

### Chemicals

All chemicals were purchased from Sigma Aldrich (USA). The polyethylene terephthalate (PET) amorphous film was purchased from Goodfellow (Germany). PET Ramapet N180 was purchased from Indorama and prepared as a semicrystalline powder by solubilisation and precipitation, as described before (Wagner-Egea et al., 2021). The crystallinity was 11.3%, determined previously by Differential Scanning Calorimetry(Wagner-Egea et al., 2023).

### Cultivation media

All chemicals used were analytical reagents. The Minimal Salt Media (MSM) and Luria-Bertani (LB) media were used in the present study. The MSM composition consists of 0.2 g KH_2_PO_4_, 1 g K_2_HPO_4_, 1 g NaCl, and 0.002 g CaCl_2_. H_2_O, 0.5 g MgSO_4_. 7H_2_O, 1 g (NH_4_)_2_SO_4_, and 0.01 g FeSO_4_.7H_2_O L^-1^. LB was prepared with 10 g tryptone, 5 g yeast extract, and 5 g NaCl L^-1^. MSM and LB agar plates were prepared by adding 16 g agar L^-1^.

### Sampling

The sample consisted of aerobic-activated sludge taken from the filtration bed of the Water filtration plant situated in Makri, Muzaffarabad AJK, Pakistan. The geographic coordinates are 34.39° of latitude and 73.47° of longitude.

### Bacterial strain isolation

The source sample was aliquoted in volumes of 10 mL, and a small piece (approx. 1.5 cm × 1 cm, and 0.02 mm thickness) of amorphous PET (Goodfellow) was submerged in each aliquot. Before use, the films were washed and sterilised with 70% ethanol. The samples were incubated at room temperature statically for 7 days. Then, the pieces of plastic were aseptically removed, washed with 0.9% NaCl, transferred to an MSM medium, and incubated for 7 days at room temperature to capture potential plastic degraders that colonised the plastic films. The growth was monitored by measuring the optical density (OD) at 600 nm wavelength. Then, the colonies were isolated by serial dilutions and spread LB agar plates.

The cultivation in a solid medium was incubated at 23°C for 48 hours. Isolated bacterial strains were transferred to LB broth with glycerol (20%) and stored at –80°C for further analysis.

### Optimization of different parameters for the growth of isolated bacteria

Inoculum size, pH, and temperature were optimised. All cultivations, carried out in triplicates, were prepared in 100 mL MSM medium containing 20 mg PET powder. The temperature range was from 23 to 43 °C, pH from 6 to 8, and inoculums of 1, 1.5, and 2 mL. All cultivations were incubated by shaking at 120 RPMs. ODs, at 600 nm wavelength, were determined every 24 hours for three days.

### Bacterial activity against PET

The ability of the isolated bacteria to degrade PET was screened by monitoring their growth in MSM medium with PET powder as the only carbon source for 28 days. All experiments were carried out in triplicate and with controls consisting of MSM medium devoid of bacteria inoculums. The cultures were prepared in 100 mL MSM containing 20 mg PET powder and incubated at 23 °C, shaking at 120 RPM. The growth was monitored spectrophotometrically by measuring OD at 600 nm.

### Growth curves

Growth curves were performed by using different carbon substrates. All cultivations were performed into 100 mL MSM containing 20 mg PET powder, 20 mg chitin, 20 mg cellulose, 1 mL tween 80, 1 mL olive oil, 200 mg glucose, or 5 mg terephthalic acid as the sole carbon source. Cultures carried out in triplicates were incubated in shake flasks at 23 °C for 24 h, and growth was monitored by measuring OD at a wavelength of 600 nm. Specific growth rates (μ_max_) and generation times were (t_g_) determined for each cultivation.

### Residual PET analysis

Residual PET from the bacterial cultivations was analysed with Fourier transform infrared (FT-IR) spectroscopy. The samples were washed with detergent (SDS 1%), rinsed with ultrapure water (milli Q water), and dried with airflow. IR spectrum was recorded in the domain of 500–4000 cm^−1^ with a 4 cm^−1^ resolution, applying 16 measured scans per sample and a data spacing of 0.964 cm^−1^. A 3.76 mA laser current was used at 30.1 °C. The spectrophotometer used was a Thermo Scientific^TM^ Nicolet iS5, and the data was analysed in the OMNIC and Origin 6.0 programs.

### Extracellular enzymes recovery

The cells were cultured in 200 mL of MSM with 20 mg of PET powder as a substrate at 23 °C for 15 days. Subsequently, the cell pellet and residual PET were separated by centrifugation at 4500 g for 20 minutes. The extracellular fraction was concentrated using spin concentration tubes with 10kD cut-off membranes. Total protein concentration was determined with the bicinchonic acid assay kit (BCA), according to the protocol provided by the manufacturer (Sigma Aldrich).

### Enzyme activity assay

Concentrated extracellular fractions were subjected to enzyme activity assay on PET powder. The semicrystalline polymer powder was soaked in 1 mL of the extracellular fraction, and the reaction mixture was incubated at 23 °C for 72 h, shaking at 200 RPM. Then, the depolymerisation products were analysed by HPLC -UV/MS.

### Analysis

The products resulting from enzymatic degradation were analysed using the HPLC-UV/MS system (Aristizábal-Lanza et al., 2022). The HPLC setup employed was the Dionex Ultimate 3000 RS, equipped with a UV/Vis detector (SPD-20A). The LC-MS system used was the ThermoFisher Scientific Ultimate 3000 RS HPLC-system, integrated with an LTQ Velos Pro Ion trap mass spectrometer and a heated electrospray ionisation source (HESI-II). The UV analysis was performed at a wavelength of 260 nm, while the mass spectrometer operated in negative mode, covering a mass range from m/z 50 to 700. The HESI source was set at 300°C with a spray voltage of 3.0 kV. Helium served as the collision gas, and nitrogen as the nebulising gas. For LC-MS/MS, data-dependent acquisition (DDA) was used with a CID fragmentation energy setting of 35%. Chromatographic separation was achieved using a hydrophobic C18 column (Kinetex® 1.7 μm XB-C18, 100 Å, LC Column 50 × 2.1 mm) with a mobile phase consisting of a 20% acetonitrile and 80% formic acid (0.1%) mixture at a constant flow rate of 0.4 ml/min for 3 minutes.

### 16S rRNA gene sequencing and phylogenetic assignment

Genomic DNA was extracted using the GeneJET™ Genomic DNA Purification Kit (Thermo Scientific). The 16S gene was PCR amplified using the universal primers 16S-FD1 (5’ GAC TTT GAT CCT GGC TCA-3’) and 16S-RP2 (5’ ACG GCT AAC TTG TTA CGA CT-3’).

PCR reactions were performed with iPoof^TM^ HF kit (Bio-Rad, USA) using 50 ng of genomic DNA as a template in 20 μL of the reaction mix. The amplification consisted of 34 cycles after an initial DNA denaturation at 95°C for 3 min and before a final polymerization at 72 °C for 5 min. Each cycle included denaturation at 95 °C for 30 s, annealing at 55 °C for 30 s, and extension at 72 °C for 1 min. The products were purified with GeneJET PCR Purification Kit #K0702 (Thermo Scientific, USA), and their quality and concentration were evaluated by agarose electrophoresis and UV spectroscopy, respectively. The amplicons were sequenced using the services of Eurofins, Germany.

The isolated bacterial strain I-MWF was identified based on a 16S rRNA gene sequence subjected to BLAST search using the Ez-Taxon server (http://eztaxon-e.ezbiocloud.net) and BLAST search on DDBJ / NCBI servers. Sequences of closely related validly published type strains used for constructing the phylogenetic tree of *Acinetobacter* strains were selected and retrieved from the EzTaxon Server (http://eztaxon-e.ezbiocloud.net) database. The phylogenetic analysis was performed with all the closely related taxa using the neighbour-joining method with the Tamura and Nei substitution model for nucleotide sequences (Saitou and Nei, 1987). Evolutionary analyses were conducted in MEGA MEGAX v10.2.6 software using the ClustalW program (Kumar et al., 2018), and the dataset was bootstrapped to 1000 iterations (Felsenstein, 1985).

### Genome sequencing, assembly, annotation, phylogenomics and comparative genomics analysis

Whole genome DNA sequencing of *A. guillouiae* I-MWF was performed GS using an Illumina NovaSeq PE150 platform (Illumina Inc.) at Novogene (China). Quality control of sequence reads was first analysed using the FastQC tool version 0.10.0 (Wingett and Andrews, 2018). Then, the reads were trimmed using the Trimmomatic program (Bolger et al., 2014). The reads assembling was performed using the SPAdes v.3.13 program (Bankevich et al., 2012). DNA contamination and confidence estimation for single-cell genome data were checked using ACDC software 2021–2022 (Lux et al., 2016). Genome annotation was predicted using online databases, including NCBI Prokaryotic Genome Annotation Pipeline (PGAP) (Tatusova et al., 2016), Clusters of Orthologous Groups (COG) (www.ncbi.nlm.nih.gov/research/cog-project) and Rapid Annotation using Subsystems Technology (RAST) (http://rast.nmpdr.org/rast.cgi). tRNA and rRNA genes were predicted by tRNAScan-SE software (Lowe and Eddy, 1997) and RNAmmer, respectively (Lagesen et al., 2007). The entire Genome Shotgun project has been deposited at DDBJ/ENA/GenBank under the accession JAMBAR000000000.1. The version described in this paper is JAMBAR010000000. A circular map of the genome *Acinetobacter* sp. strain I-MWF was constructed using CGView (Grant and Stothard, 2008).

Whole-genome-based taxonomic analysis of *A. guillouiae* I-MWF was done using the Type Strain Genome Server (TYGS) (Meier-Kolthoff and Göker, 2019). The phylogenomic tree was constructed using FastME from the Genome BLAST Distance Phylogeny (GBDP) (Lefort et al., 2015). Pair-wise-genome comparisons were conducted using GBDP and inter- genomic distances (Meier-Kolthoff and Göker, 2019). Finally, trees were rooted at the midpoint and branch supports were inferred from 100 pseudo-bootstrap replicates (Farris, 1972). OrthoVenn2 (Xu et al., 2019) was used to identify all orthologous proteins between the closest species based on the DIAMON algorithm (Buchfink et al., 2015; Wang et al., 2015). The orthologous clusters were identified using the default parameters, with a 1e-5 e-value cut- off for all protein similarity comparisons.

The comparative genomic analysis was performed on the average nucleotide identity using both best hits (one-way ANI) and reciprocal best hits (two-way ANI) between the genome sequence of *A. guillouiae* I-MWF and closest relatives identified with Whole-genome-based taxonomic analysis using the ANI calculator with the program JspeciesWS (Richter et al., 2016). Typically, the ANI values between genomes of the same species are above 95% (Rodriguez-R and Konstantinidis, 2014).

### Molecular modelling

Extracellular hydrolases with the potential to play a role in PET depolymerization were modelled using homology modelling. All annotated hydrolases from *A. guillouiae* I-MWF were subjected to signal peptide analysis using SignalP 6.0 (Teufel et al., 2022), and only those predicted to be extracellular were selected. Subsequently, the selected sequences had their signal peptides removed and were subjected to 3D modelling using the Phyre2 server (Kelley et al., 2015). Docking of MCT9977042 with (MHET)_4_ was performed previously (Wang et al., 2020a). Model analyses and figures were generated using Chimera 1.17.3 (Pettersen et al., 2004). Molecular weights and isoelectric points (pI) of the proteins were computed with ProtParam in the ExPASy server (Gasteiger et al., 2005).

## Results

### Strain identification

The isolated bacterial strain was identified as *A. guillouiae* through a comprehensive analysis of its complete 16S rDNA gene sequence. The results establish a remarkable 99.93% similarity between the isolated *A. guillouiae* strain I-MWF (MZ519889) and the reference *A. guillouiae* strain ATCC11171 (APOS01000028). *A. guillouiae*, recognized as a gram-negative, strictly aerobic bacterium, was initially isolated from gasworks effluent (Nemec et al., 2010). Within the taxonomic hierarchy, it belongs to the Pseudomonadales Order and Moraxellaceae Family, comprising 108 species (Parte et al., 2020).

Furthermore, the investigation into phylogenetic relationships based on almost-complete 16S rDNA gene sequences reveals the closest kinship of I-MWF (MZ519889) with *A. guillouiae* strain ATCC 11171 (APOS01000028), showcasing an astonishing 99.93% similarity (Figure 1). Additionally, a whole-genome-based phylogenetic tree constructed using TYGS consistently places strain I-MWF close to the *A. guillouiae* strain ATCC 11171 species (APOS01000028).

**Figure 1.**
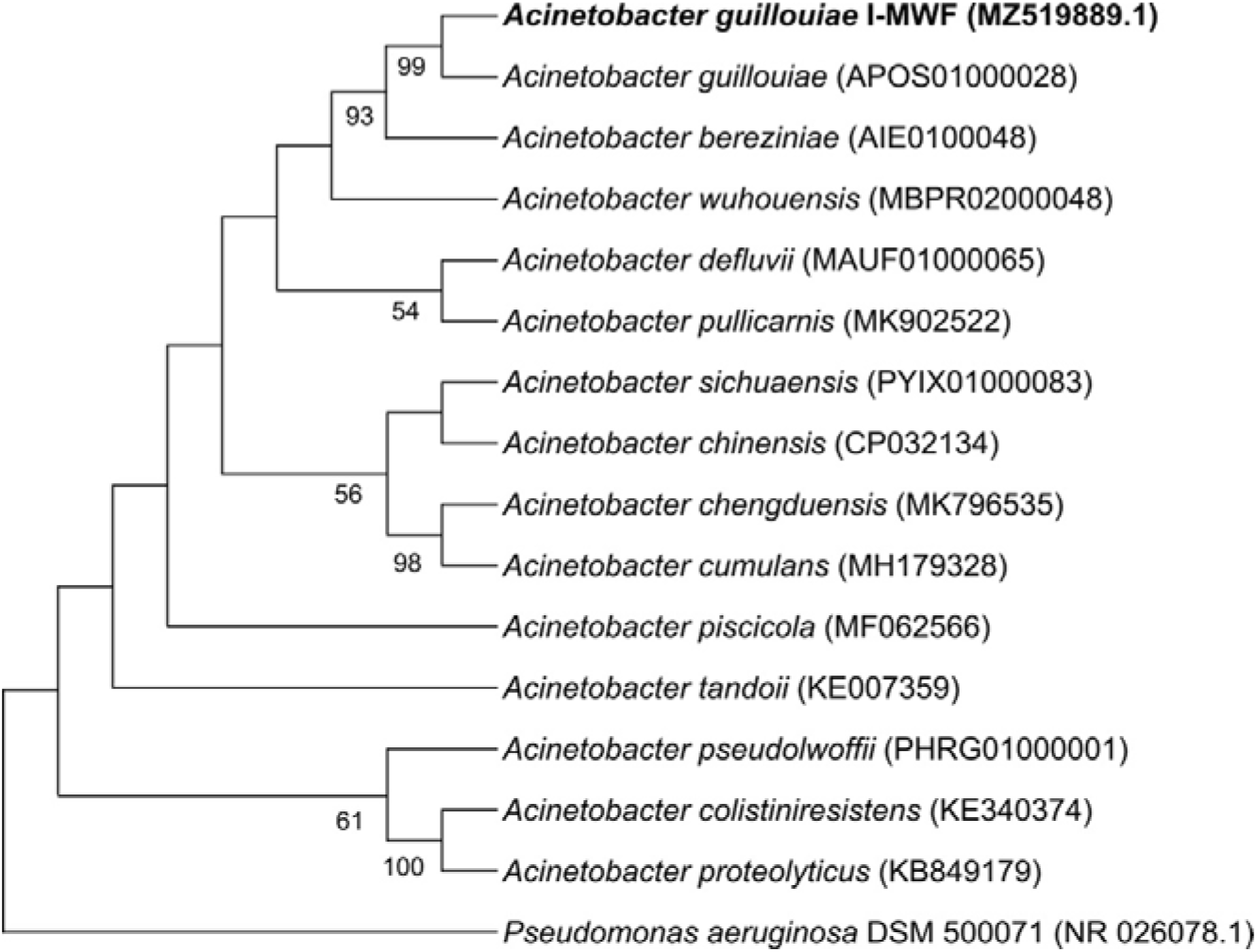
Phylogenetic tree of *Acinetobacter guillouiae* I-MWF and closest relative species based on 16S rRNA sequences. The evolutionary history was inferred using the neighbour-joining method. The percentage of replicate trees in which the associated taxa clustered together in the bootstrap test (1000 replicates) are shown next to the branches. The tree was rooted with *Pseudomonas aeruginosa* DSM (NR 026078.1). Evolutionary analyses were conducted in MEGAX v10.2.6. GenBank Accession numbers of sequences are shown in parenthesis.

To further support the previous results, we conducted an average nucleotide identity (ANI) analysis of closely related species, which indicates that the *A. guillouiae* I-MWF genome exhibits a remarkable similarity of 97% to that of *A. guillouiae* strain ATCC 11171. Significantly, this ANI value surpasses the commonly accepted 95% cutoff for species delineation (Meier-Kolthoff and Göker, 2019), thereby confirming the close relationship between the I-MWF strain and the *A. guillouiae* species.

### Isolation and growth of A. guillouiae I-MWF

The isolating strategy for *A. guillouiae* I-MWF consisted of amorphous PET film colonization followed by cultivation in MSM medium containing PET powder. PET powder was prepared by solubilization and precipitation, giving a polymer with 14 % crystallinity. The optimal temperature and pH were 23°C and 7.5, but the strain could grow in a range of pH from 7 to 8.

The growth of *A. guillouiae* I-MWF on PET powder was compared with other carbon sources after 24 hours. No growth was detected in polysaccharides such as cellulose and chitin. The highest growth was in the tween 80, followed by powder PET and olive oil, while minimal growth was detected in terephthalic acid and glucose. For deeper investigation, curve growths were performed (Figure 2), and the corresponding maximum specific growth rates and generation times were obtained (Table 1).

**Figure 2.**
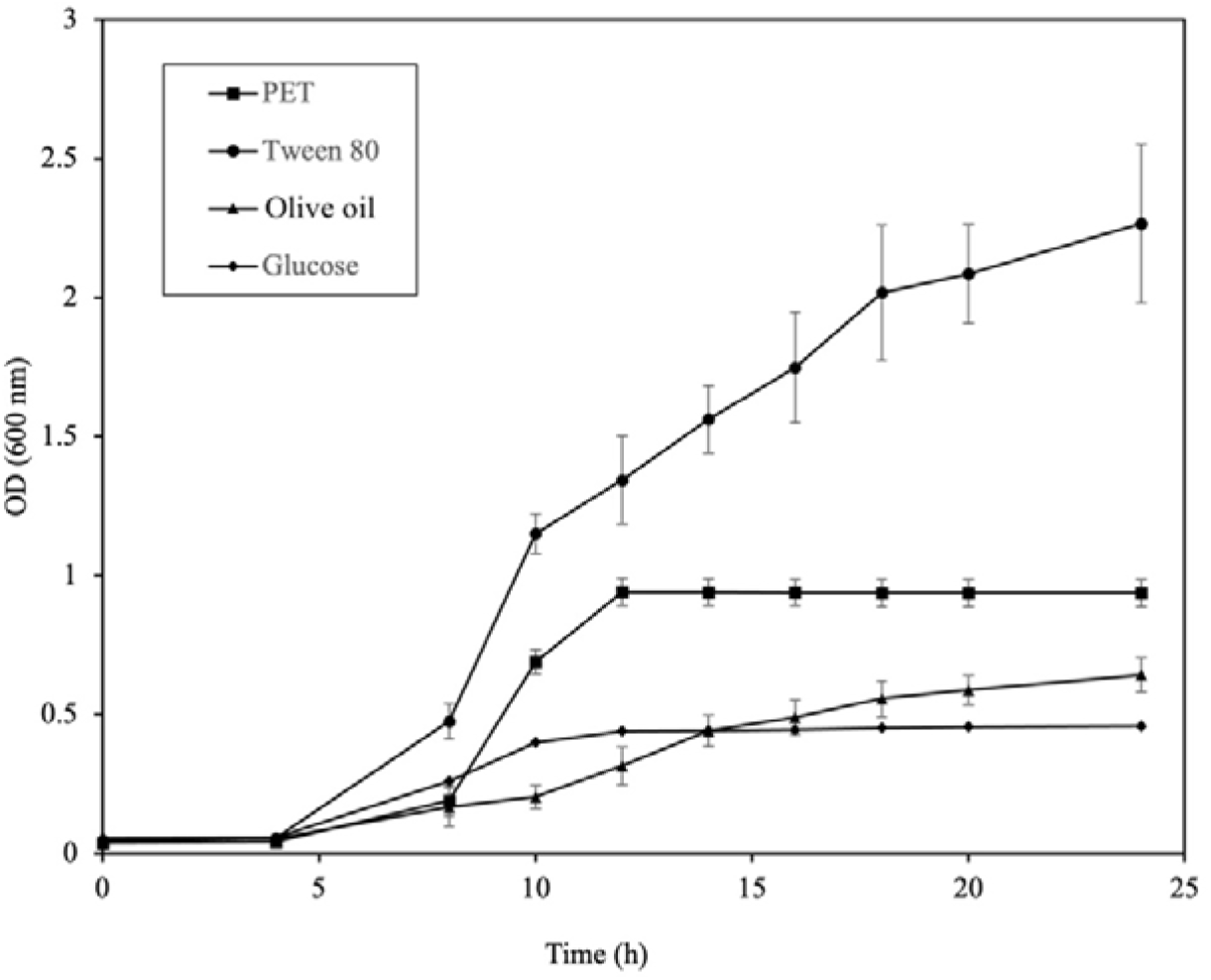
Curve growths of *A. guillouiae* I-MWF in MSM medium with different carbon sources, at 23°C.

**Table 1.** Maximum cell density (OD_max_) and kinetic parameters (μ_max_ and t_g_) for *A. guillouiae* I-MWF cultured in MSM medium with different carbon sources for 24 h at 23°C.

| Substrate | $OD_{max}$ | $\mu_{max} (h^{-1})$ | $t_g (h)$ |
| --- | --- | --- | --- |
| PET powder | 0.818 | 0.32 | 2.20 |
| Tween 80 | 2.149 | 0.38 | 1.82 |
| Olive oil | 0.59 | 0.4 | 1.73 |
| D-glucose | 0.388 | 0.23 | 3.01 |
| Terephthalic acid | 0.36 | 0.46 | 1.51 |

Residual PET powder used in the cultivation of *A. guillouiae* I-MWF was analysed by FT-IR spectroscopy to evaluate chemical changes (Figure 3). After 15 days, two new peaks appeared at 3359.89 cm^-1^ and 1644.35 cm^-1^, corresponding to O-H stretch and C=C stretching vibrations, respectively. After a month, these peaks become more intense and clearer, indicating that PET underwent aerobic microbial degradation. In particular, the peak at 1644.35 cm^-1^, ν=(C=C), became pronounced over time. This peak’s appearance is likely because of the removal of H and the formation of the double bond structure of an alkenyl group. Similar functional group changes in the backbone structure of PET by microbial degradation were also reported previously (Donelli et al., 2010). The appearance of a typical OH band owing to the intermolecular hydrogen bonding over one month is a clear indication that PET underwent hydrolysis.

**Figure 3.**
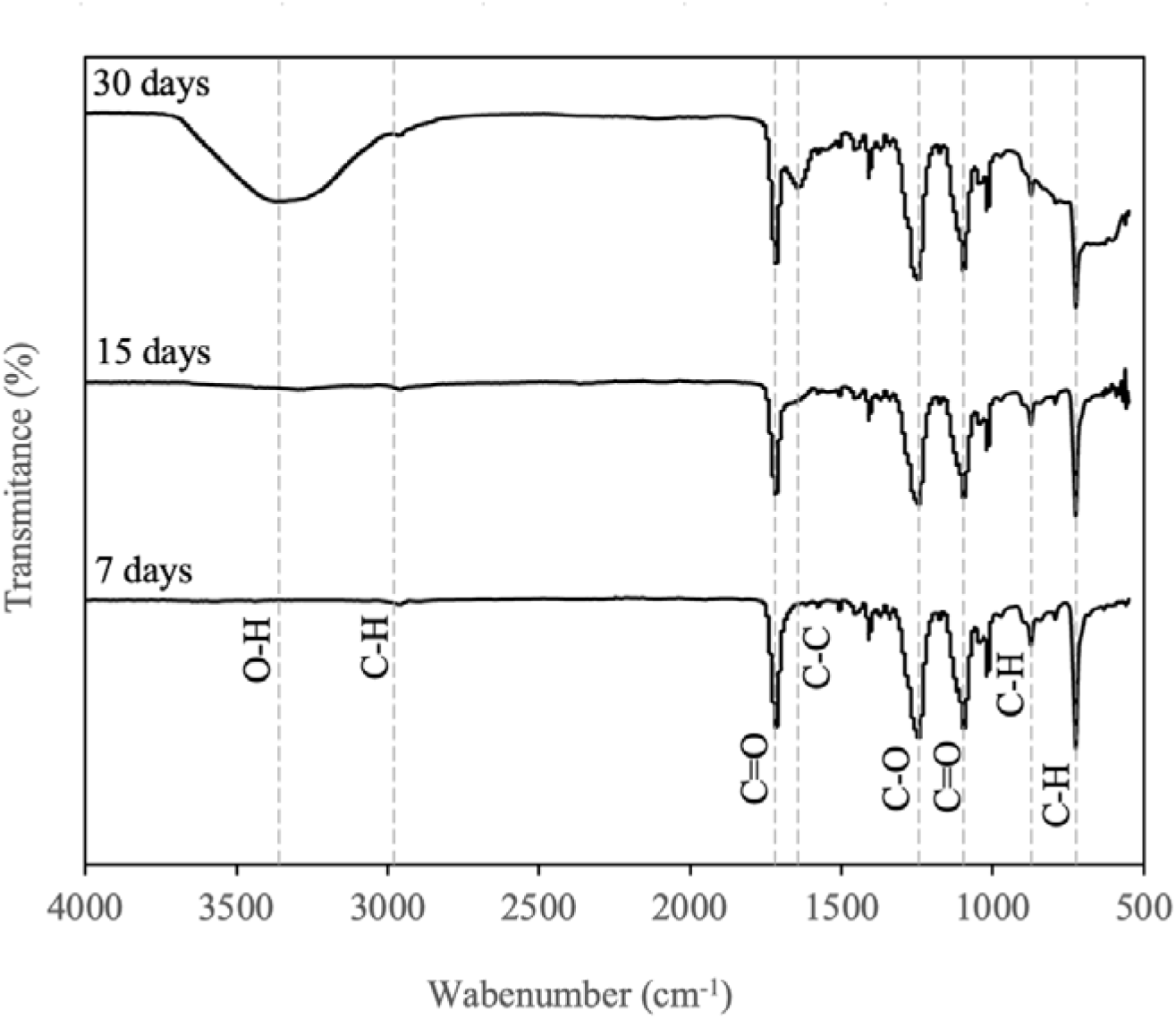
FT-IR analysis of the residual PET powder used in the *A. guillouiae* I-MWF aerobic cultivation.

### PET active enzyme analyses

The presence of extracellular PET active enzymes was determined by soaking PET powder in the extracellular medium and incubating the reaction mixture at 23 °C. TPA, MHET, and BHET were detected after 72 h of reaction by LC-MS analysis (Figure 4A). When the reaction was monitored for a longer time, it was observed that TPA and MHET were produced first and increased with time. In the sample analysed after 30 days, BHET was also quantifiable (Figure 4B). The highest concentration of depolymerisation products was 86 mg/L TPA, 159 mg/L MHET, and 2 mg/L BHET. Although easily detectable, the low concentrations of products could reflect the low content of PET active enzymes in the extracellular samples and enzyme inactivation over time. The total protein was 0.15±0.02 mg/mL. However, the PET active enzymes likely represent only a small fraction of them.

**Figure 4.**
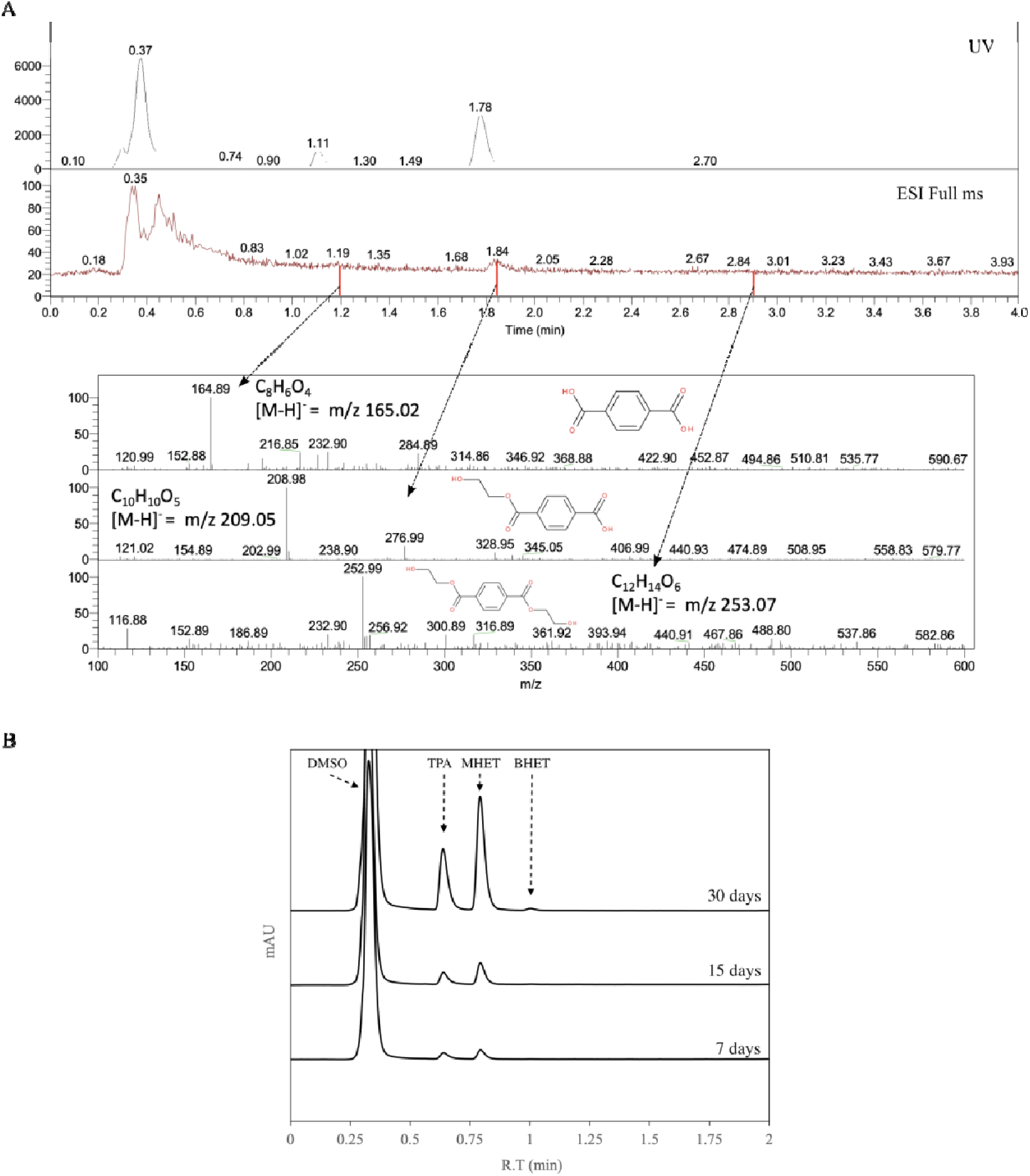
Analysis of PET depolymerization products in the extracellular medium of *A. guillouiae* I-MWF. (A) LC-MS analysis. (B) HPLC-UV analysis.

### Genome features

The *A. guillouiae* I-MWF genome comprises 87 scaffolds of 4.61 Mb total, with an average GC content of 38.2% and N50 of 4.6 Mb (Table 2). The NCBI Prokaryotic genome annotation pipeline (PGAP) feature annotation produced a total of 4,303 genes, from which 4,178 were identified as protein-coding genes, along with six rRNA (three 5S, two 16S, one 23S) and 71 tRNA genes (Table 2, Figure 5). Genomes of other strains of *A. guillouiae* species (n=16) deposited in the GenBank database range from 4.91Mb (*A. guillouiae* NIPH 991; APPJ01) to 4.55 Mb (*A. guillouiae;* JAHPTS01) (Table 3).

**Figure 5.**
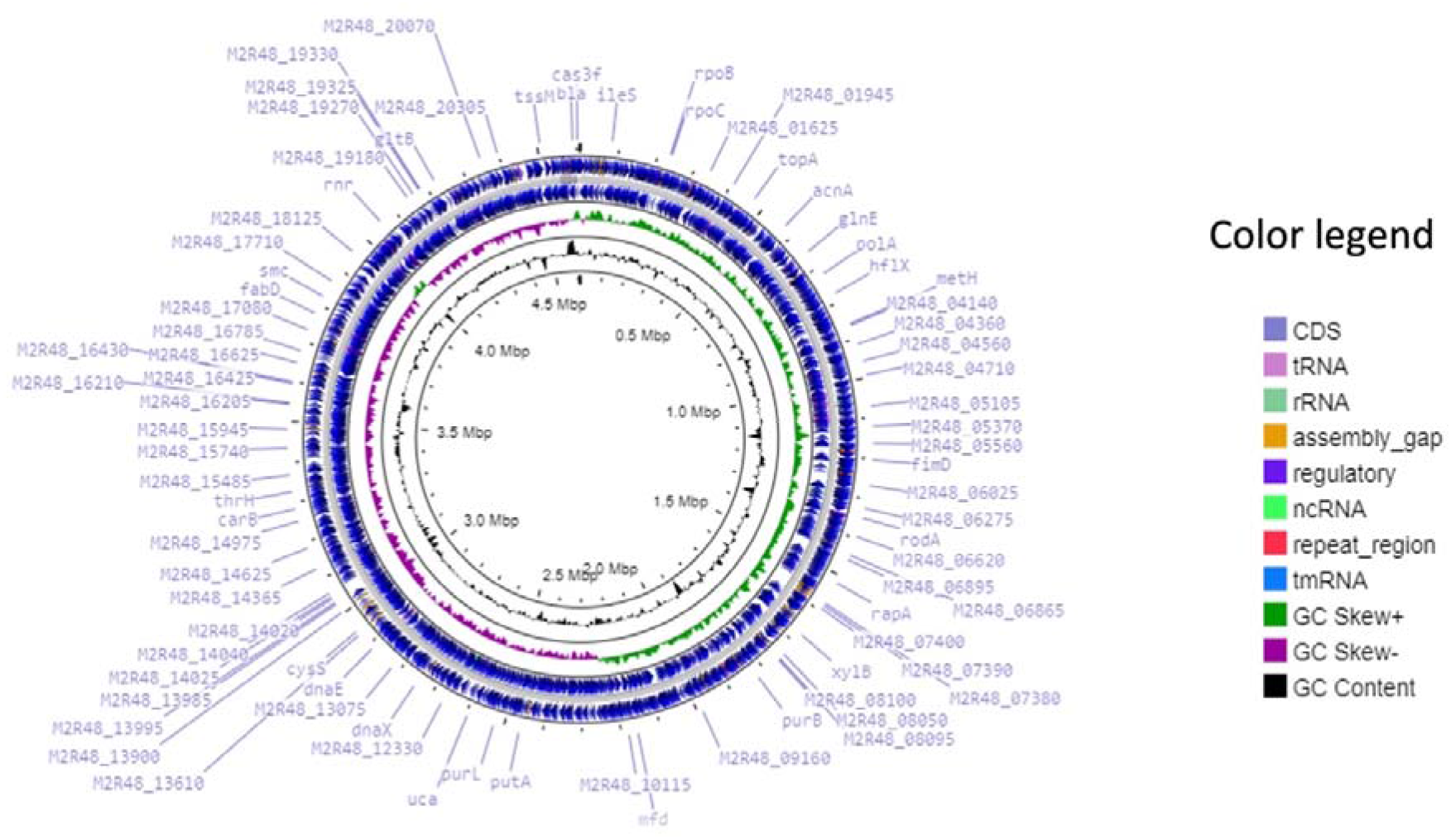
Graphical circular genome map of *A. guillouiae* I-MWF genome constructed using CGView (http://cgview.ca). The outermost ring shows the coding sequences with PGPA annotation (including tRNA, rRNA, tmRNA and ncRNA). The GC content is represented in black. The GC skew pattern is represented by the innermost ring, with purple indicating negative values and green indicating positive values.

**Table 2.** General features of the *A. guillouiae* I-MWF genome.

| Item | Value |
| --- | --- |
| Number of scaffolds | 87 |
| Total contig length (Mb) | 4.61 |
| G + C content (%) | 38.2 |
| Number of protein coding genes | 4,178 |
| Number of predicted transfers RNAs | 71 |
| Number of predicted ribosomal RNAs | 6 |
| GenBank accession number | JAMBAR000000000.1 |

**Table 3.** General genomic features of *Acinetobacter guillouiae* species deposited in GenBank database (as of August 2023).

| Strain | Level | Size (Mb) | GC% | WGS | Scaffolds | CDS |
| --- | --- | --- | --- | --- | --- | --- |
| CC160 | Contig | 4.64321 | 38.1 | JAGIPP01 | 16 | 4123 |
| SFC 500-1A | Contig | 4.81476 | 38.1 | JAHWXT01 | 41 | 4346 |
| BIGb0206 | Contig | 4.67876 | 38 | JANUEF01 | 31 | 4291 |
| WU_MDCL_Ag157 | Contig | 4.55717 | 38.1 | JAHPTS01 | 49 | 4152 |
| KCTC 23200 | Contig | 4.84278 | 38.1 | BBRY01 | 72 | 4486 |
| UN03-107 | Scaffold | 4.70491 | 38.2 | JAQEOA01 | 85 | 4292 |
| UN03-29 | Scaffold | 4.69634 | 38.2 | JAQEPJ01 | 88 | 4300 |
| AM113-120 | Scaffold | 4.6844 | 38.2 | JAQDVI01 | 102 | 4291 |
| TUM15313 | Contig | 4.67043 | 38.2 | BKPA01 | 182 | 4237 |
| TUM15493 | Contig | 4.74103 | 38.2 | BKVV01 | 284 | 4287 |
| PgBE136 | Chromosome | 4.6231 | 38.1 | Not available | 1 | 4144 |
| CCUG 2491 | Contig | 4.87842 | 38.1 | VZOG01 | 78 | 4515 |
| NBRC 110550 | Complete | 4.64842 | 38.3 | Not available | 1 | 4164 |
| NIPH 991 | Scaffold | 4.90577 | 38.2 | APPJ01 | 4 | 4417 |
| CIP 63.46 | Scaffold | 4.90228 | 38.2 | APOS01 | 16 | 4479 |
| MSP4-18 | Contig | 4.84896 | 38.1 | ASQG01 | 94 | 4485 |

Based on sequence homology, coding domain sequences (CDSs), we categorized 3872 genes into COGs (Clusters of Orthologous Groups) (Tatusov et al., 2000), including putative or hypothetical genes and genes of unknown function, mapped into 22 different COGs (Figure 6). The most abundant COG categories apart from the [S] function unknown (886 CDSs) and unclusterd COGs (570 CDS), were those assigned to [R] general function prediction (570 CDSs, 11.5%), [k] transcription (303CDSs, 6.11%), [E] amino acid transport and metabolism (295 CDSs, 5.95%), [P] inorganic ion transport and metabolism (258, CDSs, 5.2%), [C] energy production and conversion (219, CDSs, 4,41%), [M] cell wall/membrane biogenesis (218, CDSs, 4.39%), [J] translation (205 CDSs, 4.13%), [I] Lipid transport and metabolism (205 CDSs, 4.13%) (Figure 6).

**Figure 6.**
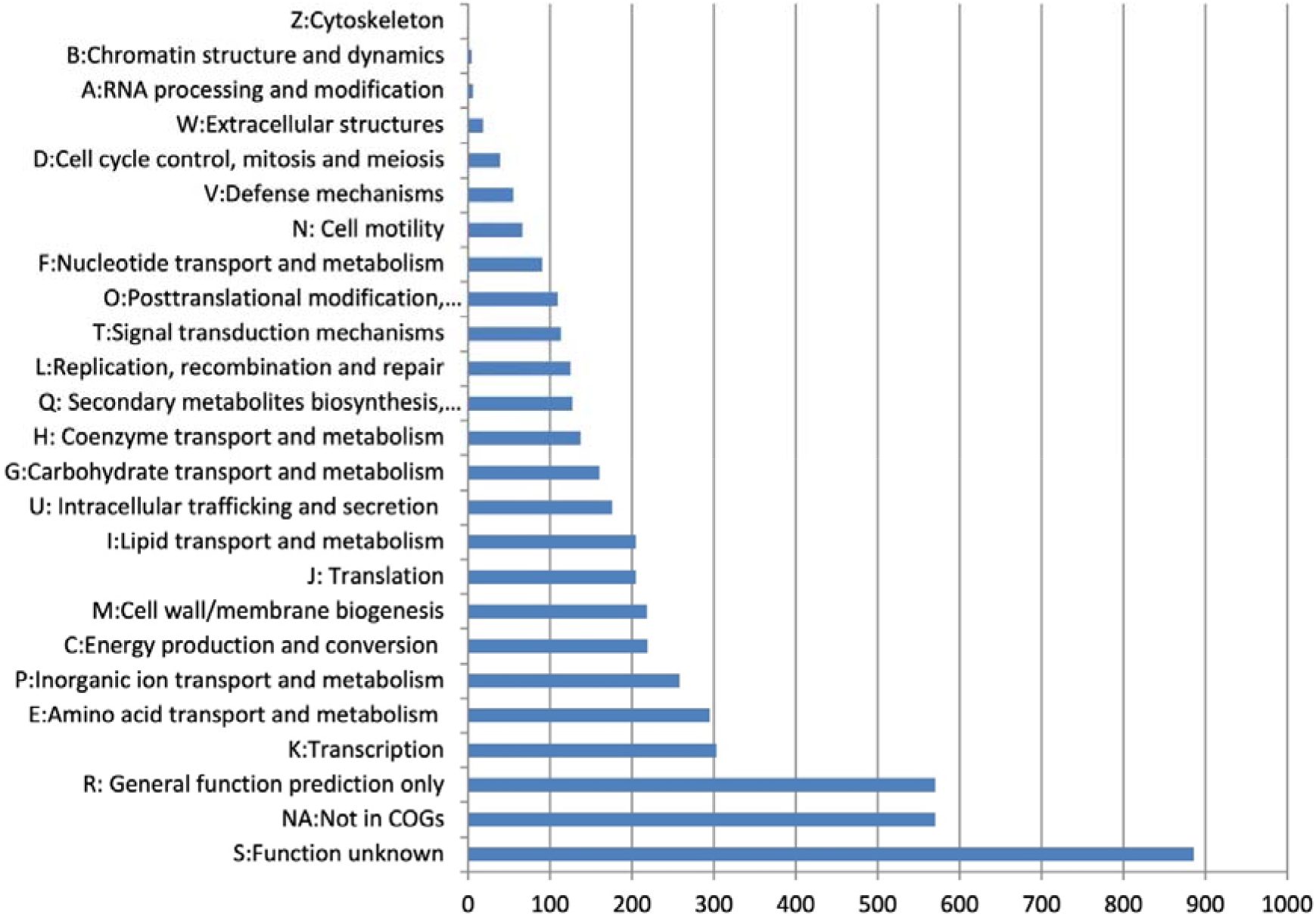
COG functional classification of *A. guillouiae* I-MWF genome. The X-axis represents the number of genes; the Y-axis is the COG function class.

The subsystem distribution and general information on the potential functional distribution based on RAST annotation are illustrated in Figure 7. Genes associated with amino acids and derivatives (387 ORFs), carbohydrates (286 ORFs), cofactors, vitamins, prosthetic groups, and pigments (257 ORFs), and protein metabolism (250 ORFs). Fatty Acids, lipids, and isoprenoids (189 ORFs) were abundant among the SEED subsystem categories.

**Figure 7.**
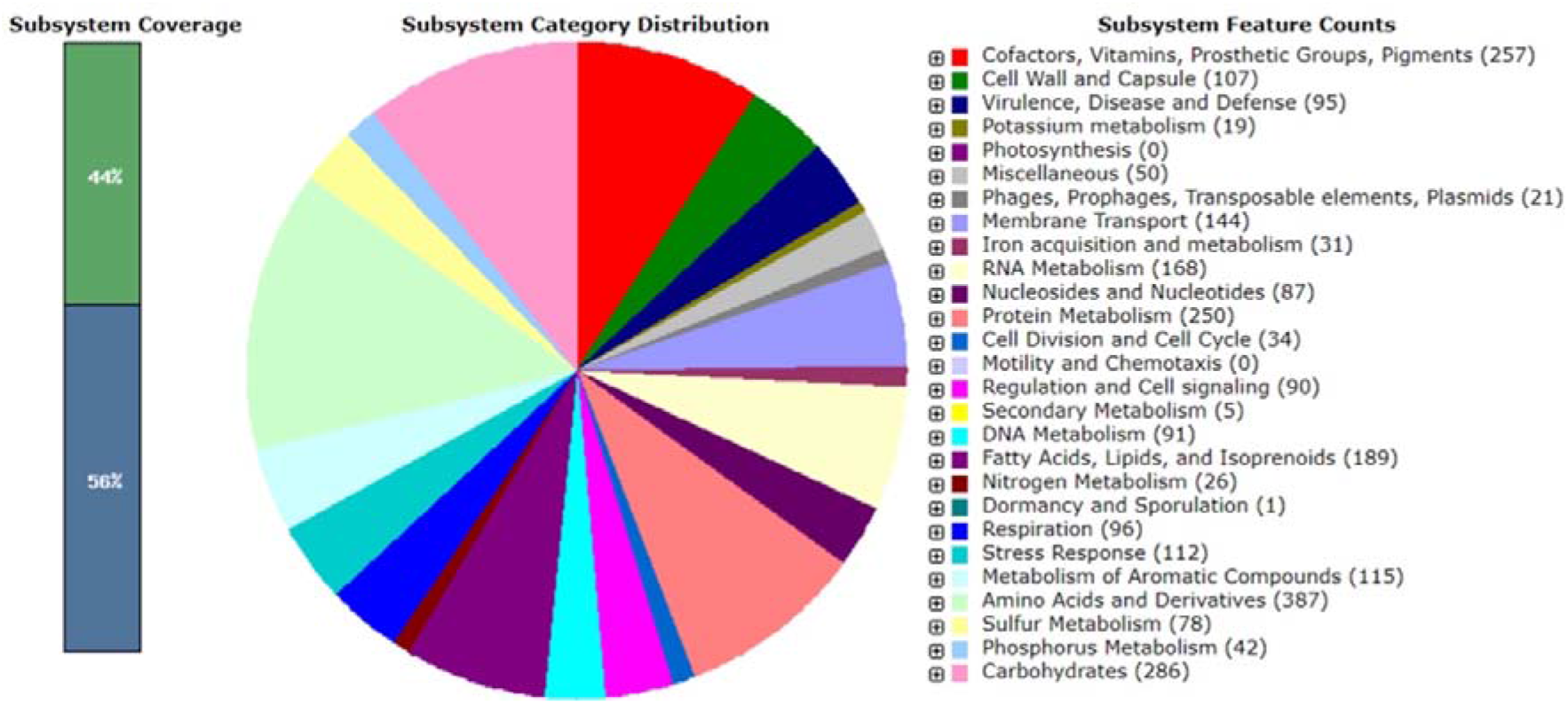
An overview of the subsystem categories assigned to the metabolic genes and pathways predicted in the genome of *A. guillouiae* I-MWF genome. The green part in the bar chart at the leftmost position corresponds to the percentage of proteins involved (44%), and the pie chart in the right panel demonstrates the fraction and count (parenthesis in legend) of each subsystem feature.

### Comparative genomic analysis

A whole-genome-based phylogenetic tree created using the TYGS server showed that strain I- MWF is close to the *A. guillouiae* CIP 63.46 (ATCC 11171) (APOS01000028), which confirms the result of 16SrDNA-based analysis (Figure 8). The ANI values of pressure I- MWF with all genomic sequences of *A. guillouiae* strains deposited in GenBank (since 2023) are more than 97%, which is above the 95% cutoff for the same species, which confirms the assignment of I-MWF strain to *A. guillouiae* species.

**Figure 8.**
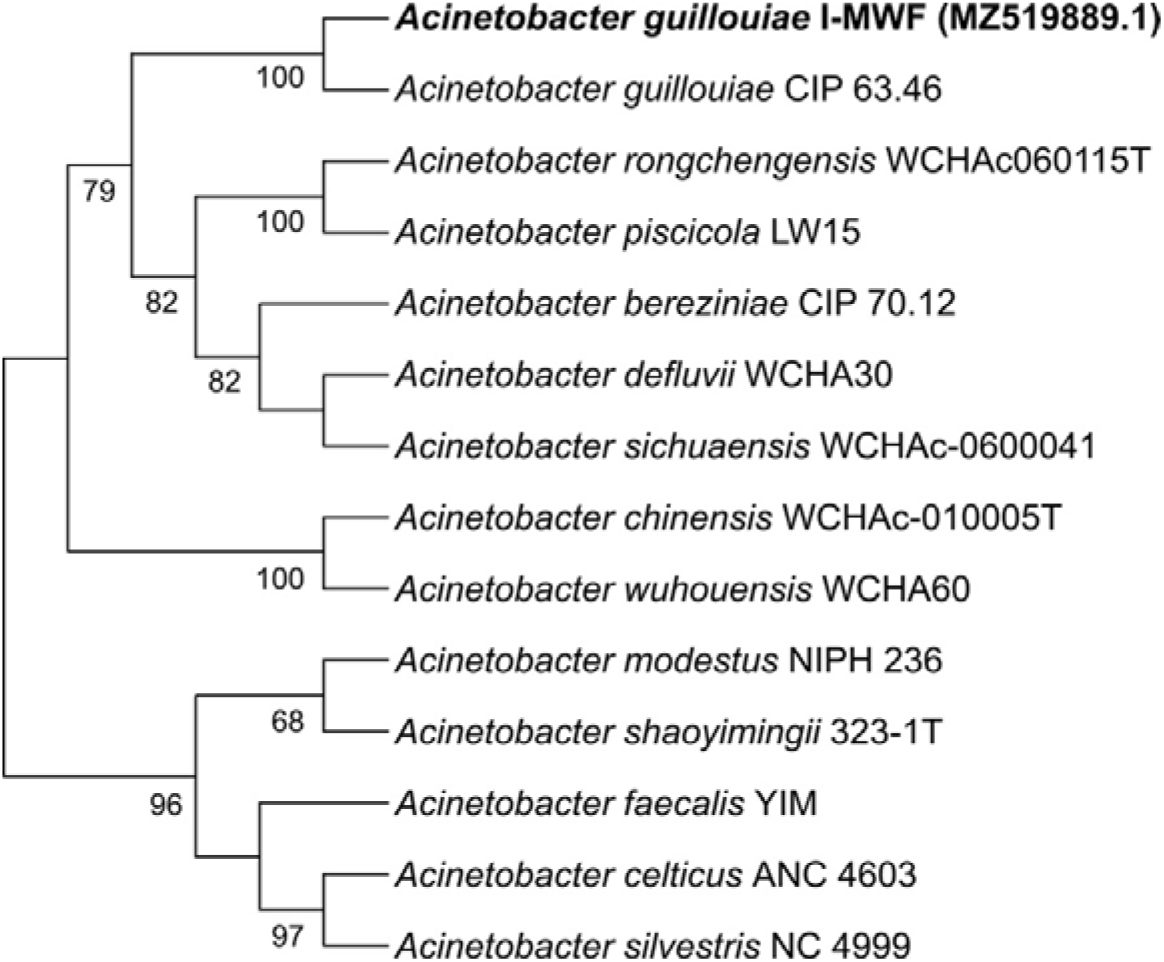
Phylogenomic tree based on genome sequences in the TYGS tree inferred with FastME 2.1.6.1 from Genome BLAST Distance Phylogeny approach (GBDP). The branch lengths are scaled in terms of the GBDP distance formula d5. The numbers above branches are GBDP pseudo-bootstrap support values>50% from 100 replications. The tree was rooted at the midpoint.

Pangenome analysis of *A. guillouiae* I-MWF with the closest species, *A. guillouiae* CIP 63.46 (ATCC 11171), was done using orthoVenn2 software (Figure 9). The pangenome consisted of 3678 clusters, with 65 orthologous clusters (clusters with at least two copies from one genome) and 3613 single-copy gene clusters. The core genome contained 3929 proteins. Strain-specific clusters were unevenly distributed; the I-MWF strain showed 40 clusters potentially assigned to P: fatty acid biosynthetic process, P: cell adhesion, F: translation elongation factor activity, P: pathogenesis, P: protein secretion by the type V secretion system, F: uridine phosphorylase activity, F: protein serine/threonine kinase activity, and P: polysaccharide catabolic process. The remaining clusters are unannotated (with no Swiss-Prot Hit or Gene Ontology Annotation). The molecular functions assigned to all these clusters are transferase activity (GO:0016740) and nucleic acid binding (GO:0003676). Concerning *A. guillouiae,* nine specific clusters were identified and assigned to P: translesion synthesis (GO:0019985), P: lipopolysaccharide biosynthetic process (GO:0009103), P: regulation of transcription (GO:0006355), F: transferase activity (GO:0016757), and the other 5 clusters remained unannotated. The molecular function assigned these clusters to transferase activity (GO:0016740). These results showed although the two strains belong to the same species, each genome encodes several unique features, which may be due to the ecological niche adaptation.

**Figure 9.**
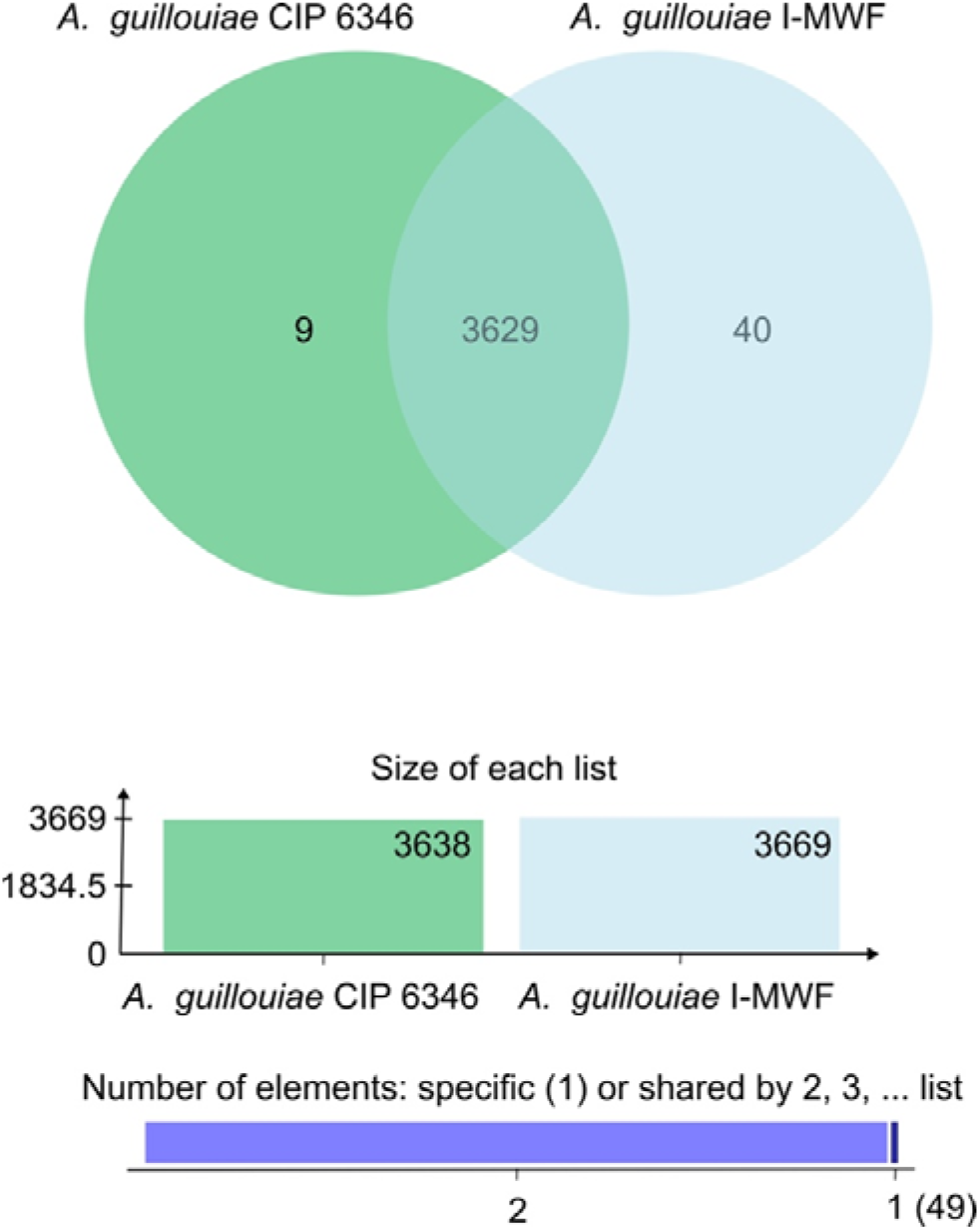
Venn diagram showing the distribution of shared and unique orthologous gene clusters among *A. guillouiae* I-MWF and *A. guillouiae* CIP 63.46 strains as visualized by OrthoVenn2. A total of 3629 shared clusters of orthologous groups were identified in these 2 strains.

### Putative enzymes involved in PET degradation

The *A. guillouiae* I-MWF genome sequence allowed the annotation of at least 18 genes encoding extracellular hydrolases. The molecular weight (MW) and isoelectric point (pI) were theoretically obtained from the amino acid sequences (Table 4). The results revealed a wide-ranging diversity in MW, spanning from 20,010 Da to 47,465 Da, with an average MW of around 34,000 Da. Similarly, pI values exhibited significant variability, ranging from 4.99 to 9.86, indicating adaptation to a broad pH spectrum. This MW and pI values diversity suggests that these extracellular proteins have diverse functions in bacterial physiology. In addition, certain proteins exhibited remarkable properties, such as the high pI of MCT9979697 and the high MW of MCT9979753.

**Table 4.**
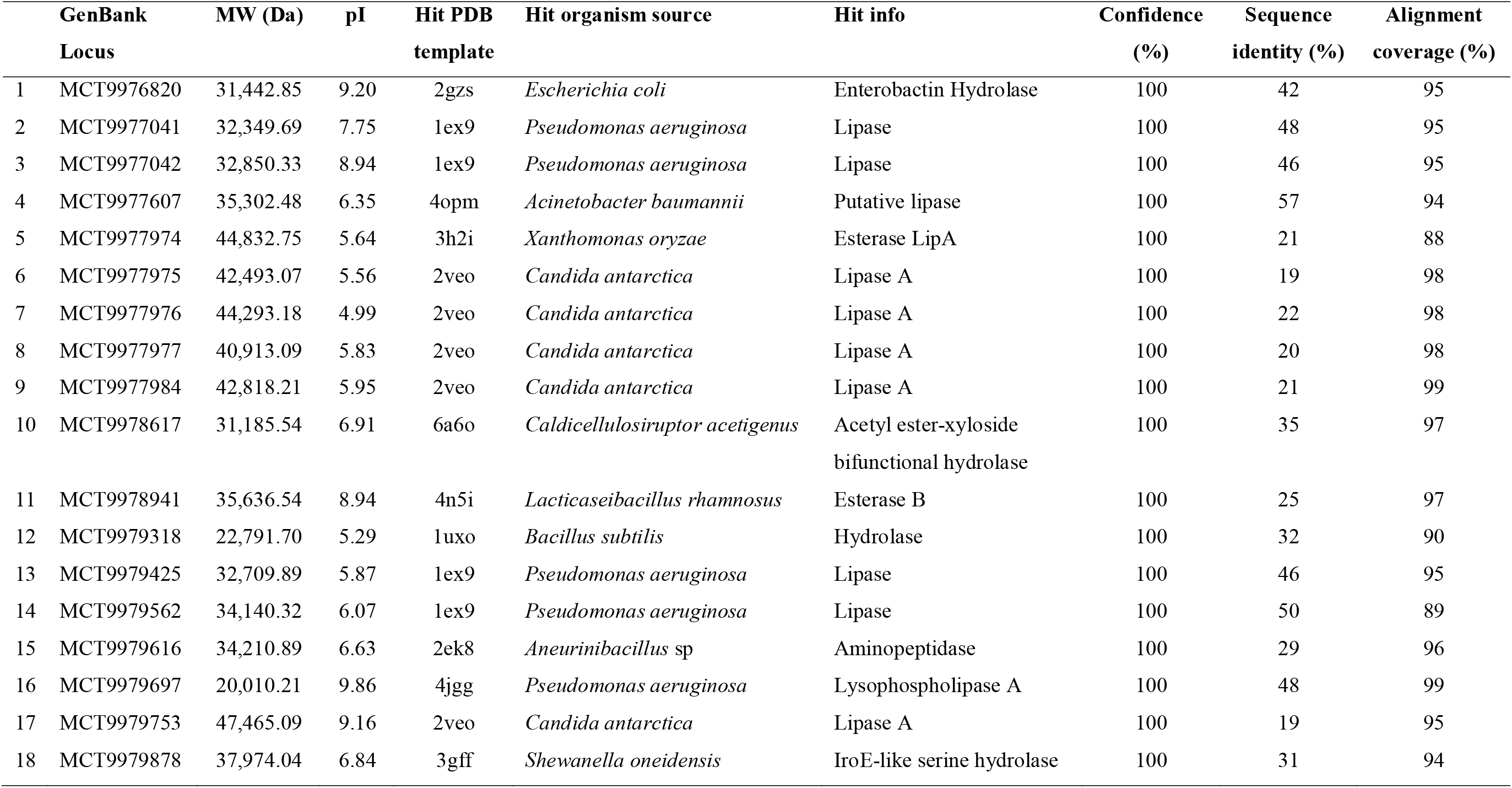
Molecular modelling of extracellular hydrolases selected from the annotated genome of *A. guillouiae* I-MWF. MW are the molecular weights, and pI are the lectric points.

|  | GenBank<br>Locus | MW (Da) | pI | Hit PDB<br>template | Hit organism source | Hit info | Confidence<br>(%) | Sequence<br>identity (%) | Alignment<br>coverage (%) |
| --- | --- | --- | --- | --- | --- | --- | --- | --- | --- |
| 1 | MCT9976820 | 31,442.85 | 9.20 | 2gzs | *Escherichia coli* | Enterobactin Hydrolase | 100 | 42 | 95 |
| 2 | MCT9977041 | 32,349.69 | 7.75 | 1ex9 | *Pseudomonas aeruginosa* | Lipase | 100 | 48 | 95 |
| 3 | MCT9977042 | 32,850.33 | 8.94 | 1ex9 | *Pseudomonas aeruginosa* | Lipase | 100 | 46 | 95 |
| 4 | MCT9977607 | 35,302.48 | 6.35 | 4opm | *Acinetobacter baumannii* | Putative lipase | 100 | 57 | 94 |
| 5 | MCT9977974 | 44,832.75 | 5.64 | 3h2i | *Xanthomonas oryzae* | Esterase LipA | 100 | 21 | 88 |
| 6 | MCT9977975 | 42,493.07 | 5.56 | 2veo | *Candida antarctica* | Lipase A | 100 | 19 | 98 |
| 7 | MCT9977976 | 44,293.18 | 4.99 | 2veo | *Candida antarctica* | Lipase A | 100 | 22 | 98 |
| 8 | MCT9977977 | 40,913.09 | 5.83 | 2veo | *Candida antarctica* | Lipase A | 100 | 20 | 98 |
| 9 | MCT9977984 | 42,818.21 | 5.95 | 2veo | *Candida antarctica* | Lipase A | 100 | 21 | 99 |
| 10 | MCT9978617 | 31,185.54 | 6.91 | 6a6o | *Caldicellulosiruptor acetigenus* | Acetyl ester-xyloside<br>bifunctional hydrolase | 100 | 35 | 97 |
| 11 | MCT9978941 | 35,636.54 | 8.94 | 4n5i | *Lactocaseibacillus rhamnosus* | Esterase B | 100 | 25 | 97 |
| 12 | MCT9979318 | 22,791.70 | 5.29 | 1uxo | *Bacillus subtilis* | Hydrolase | 100 | 32 | 90 |
| 13 | MCT9979425 | 32,709.89 | 5.87 | 1ex9 | *Pseudomonas aeruginosa* | Lipase | 100 | 46 | 95 |
| 14 | MCT9979562 | 34,140.32 | 6.07 | 1ex9 | *Pseudomonas aeruginosa* | Lipase | 100 | 50 | 89 |
| 15 | MCT9979616 | 34,210.89 | 6.63 | 2ek8 | *Aneurinibacillus* sp | Aminopeptidase | 100 | 29 | 96 |
| 16 | MCT9979697 | 20,010.21 | 9.86 | 4jgg | *Pseudomonas aeruginosa* | Lysophospholipase A | 100 | 48 | 99 |
| 17 | MCT9979753 | 47,465.09 | 9.16 | 2veo | *Candida antarctica* | Lipase A | 100 | 19 | 95 |
| 18 | MCT9979878 | 37,974.04 | 6.84 | 3gff | *Shewanella oneidensis* | IroE-like serine hydrolase | 100 | 31 | 94 |

The molecular structures of extracellular enzymes potentially exhibiting ester hydrolase activity were modelled through computational simulations using the Phire2 server, boasting a confidence level of 100% (Table 4). Regarding the crystallographic templates employed, alignment coverage ranged from a minimum of 88% to a maximum of 99%, while sequence identities spanned from 19% to 57%. It’s worth noting that although no enzymes resembling cutinases were identified, our analysis yielded a diverse array of potential lipases and three esterases. All these structures shared the typical α/β hydrolase fold.

The crystal structure of the lipase 1ex9 (PDB ID: 1ex9) from *Pseudomonas aeruginosa* has been the template for four lipases, with identities higher than 46% in covers superior to 89%. The overall structures consist of a β-sheet of parallel strands surrounded by α-helixes. All these lipases were modelled with open-form active sites (Figure 10A). The lipase A (PDB ID: 2veo) from *Candida antarctica* has been the template for five lipases with identities ranging from 19 to 22% with covers higher than 95% (Figure 10B).

**Figure 10.**
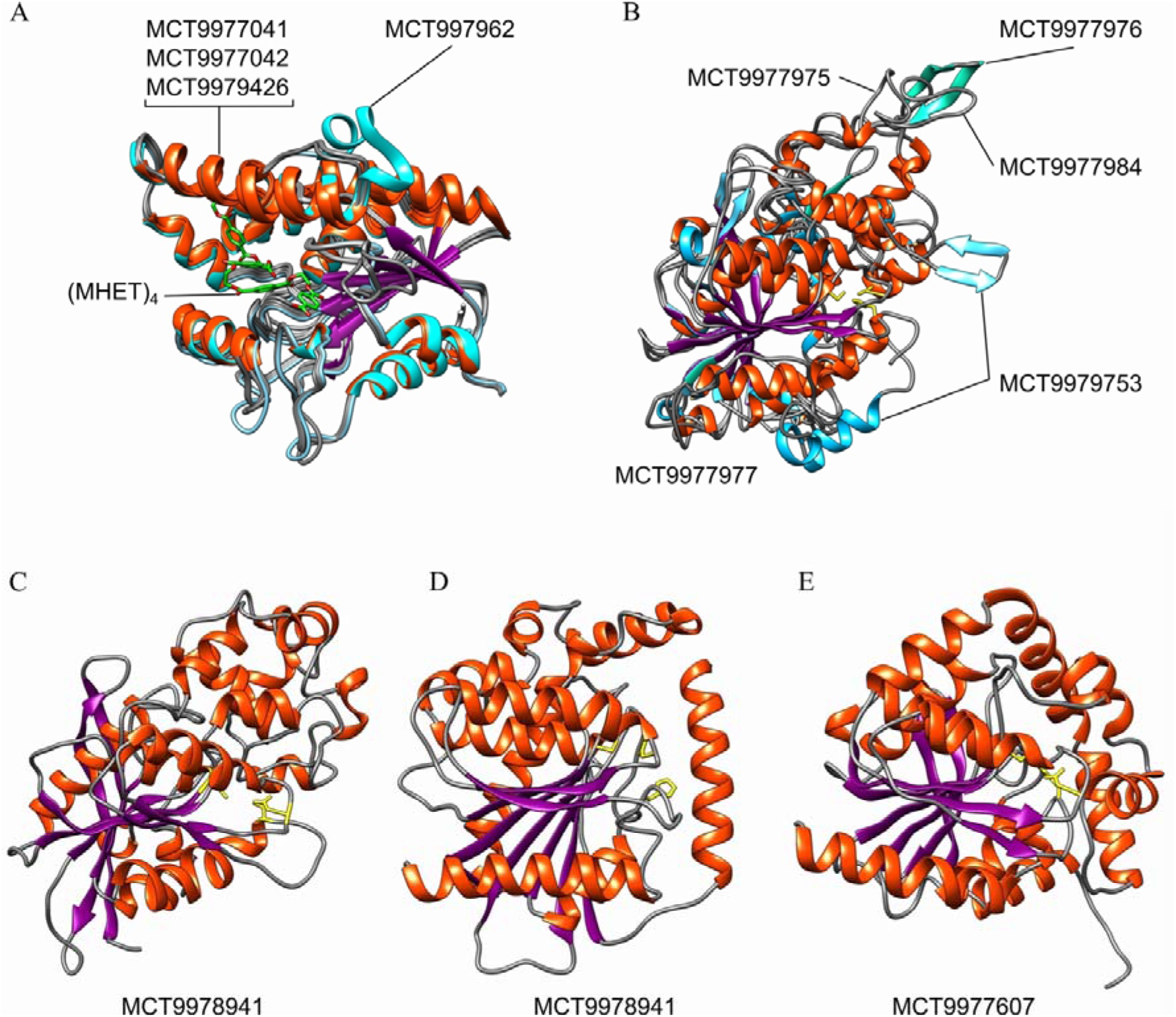
Molecular models of representative putative lipases and esterases from *A. guillouiae* I-MWF. (A) Overlapped models based on lipase 1xe9 from *Pseudomonas aeruginosa*. Docking of MCT9977042 with (MHET)_4_, the ligand is shown in green. (B) Overlapped models based on lipase A 1xe9 from *Candida antarctica*. (C) Model-based in esterase LipA h32i from *Xanthomonas oryzae*. (D). Model-based in esterase B 4n5i form *Lacticaseibacillus rhamnosus*. (E) Model-based on a putative lipase from *Acinetobacter baumannii*. In all models, the predicted catalytic triad is in yellow.

The esterase LipA (PDB ID: 3h2i) from *Xanthomonas oryzae* has been the best template, although with a low identity (21 %) in a cover of 88% (Figure 10C). Another esterase was predicted by using the esterase B (PDB ID: 4n5i) from *Lacticaseibacillus rhamnosus* as the best template, sharing 25% of identity in a cover of 97% (Figure 10D). A third esterase was predicted based on the crystal structure (PDB ID: 4opm) of a putative lipase from *Acinetobacter baumannii*, with an identity of 57% and a cover of 94% (Figure 10E).

In addition, putative proteins involved in terephthalic acid metabolism were selected based on their potential activity on phenolic compounds and their metabolic intermediates. Thus, genes encoding enzymes active on aromatic compounds, phenol, benzoate, catechol, extradiol, mucolactone, mucronate, and carboxymucolctone were found in the genome of *A. guillouiae* I-MWF. In addition, genes encoding for a benzoate H/(+) symporter and MSF transporter (Table 5).

**Table 5.**
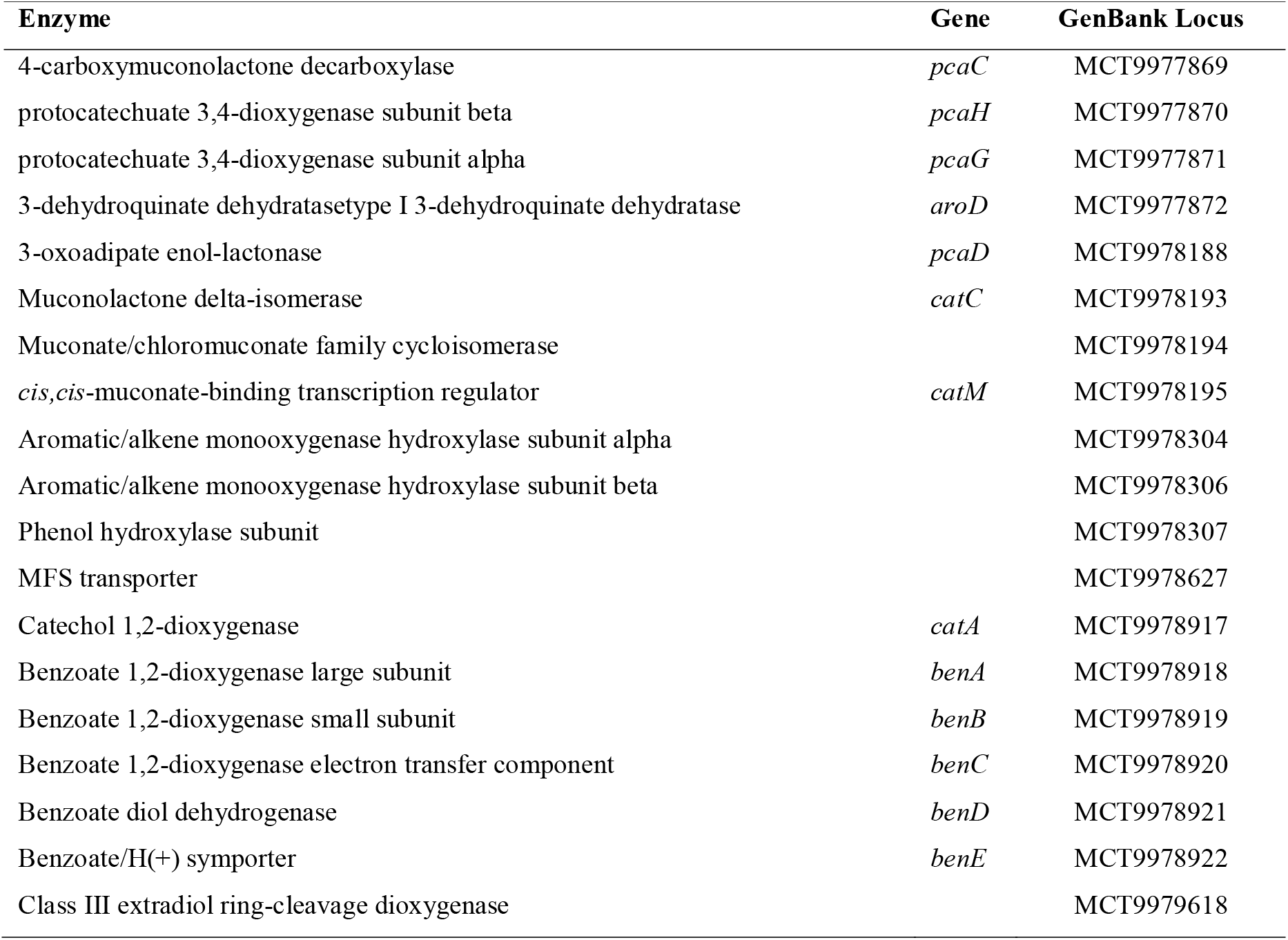
Annotated proteins involved in the metabolism of phenolic compounds in *A. guillouiae* I-MWF.

## Discussion

Sludge is a source of inorganic and organic nutrients that facilitate microbial growth, including microorganisms with potential PET degradation activity (Torena et al., 2021). In the present work, *A. guillouiae* I-MWF was successfully isolated by introducing an amorphous PET film into sludge samples to select bacteria capable of colonizing the plastic and subsequent strain isolation in enrichment culturing. For this strategy, we assumed that film- colonizing bacteria could develop the ability to depolymerize PET. In addition, amorphous PET films could also be more accessible to bacterial attack than highly crystalline ones, facilitating their colonization.

*A. guillouiae* I-MWF has initially been identified by phylogenetic analysis of the complete ribosomal gene 16S. A phylogenomic analysis and comparative genomics have corroborated the species. The 16S phylogeny shows that *A. guillouiae* I-MWF is closely related to *A. bereziniae* (Figure 1). The genomic analysis revealed that the strain *A. guillouiae* CIP 63.44 is the closest to *A. guillouiae* I-MWF (Figures 8 and 9), although they have unique features attributed to ecological niche adaptations. *A. guillouiae* strains have been found mainly in sludge. However, recently, an *A. guillouiae* strain able to degrade polyethylene by alkane hydroxylation and alcohol dehydrogenation has been isolated from insect larvae (Kim et al., 2023).

Several *Acinotebacter* species attracted attention for their importance in bioremediation. For example, *A. calcoaceticus* PA, isolated from petrochemical wastewater, degrades phenol (Liu et al., 2016). *A. tandoii* isolated from the gut of termite also degrades phenol (Van Dexter and Boopathy, 2019). *A. caloaceticus* could degrade bisphenol A (Kwon et al., 2006). *Acinetobacter* sp. DSM 17874 could grow using n-alkanes with chains ranging from C10 to C40 as a sole carbon source (Throne-Holst et al., 2007). *Acinetobacter baumannii* DP-2 degraded dibutyl phthalate (Li et al., 2022)*. A. oleivorans* could grow in diesel oil (Kang et al., 2011). A newly isolated *Acinetobacter* sp. strain LMB-5 was able to degrade phthalate esters (Yue et al., 2017). Also, some species can degrade polymers. An *Acinetobacter* sp. strain isolated from the *Tribolium castaneum* larvae was associated with polystyrene degradation (Wang et al., 2020b). Another *Acinetobacter* sp. strain found in the *Tenebrio molitor* larvae could degrade polyethylene films (Yin et al., 2020). *Acinetobacter pittobacter* showed a degradation effect on low-density polyethylene (Zhang et al., 2023). Thus, these reports show that various *Acinetobacter* species are equipped with enzymatic machinery able to metabolize both aromatic and aliphatic hydrocarbons.

In this study, we found, to the best of our knowledge, for the first time, that an *A. guillouiae* strain can grow on and partially degrade PET powder (Figure 2, Table 1). Furthermore, the growth of *A. guillouiae* I-MWF in Tween 80 and olive oil shows its ability to utilize lipoic substances. In contrast, it could not grow on carbohydrates such as cellulose and chitin and only showed minimal growth on glucose. The highest maximum growth rate and cell density were achieved with Tween 80 as carbon source, μ_max_ 0.32 h^-1^, and OD_600nm_=2.149 in 24 h, respectively. Although *A. guillouiae* I-MWF is partially grown on PET powder, there is no evidence of using PET as the sole carbon source. Residual yeast extract from the inoculum could provide a carbon source for partial growth. In the cultivation medium containing PET powder, the maximum growth rate was 0.32 h^-1^, close to that obtained in Tween 80, but with a significantly lower maximum cell density, reaching the stationary phase at OD_600nm_=0.818, indicating the depletion of the available carbon source despite sufficient PET in the culture medium. This could be attributed to the fact that *A. guillouiae* I-MWF ceased growing when residual yeast extract was depleted. In addition, the strain could not grow in non-pre-treated PET, such as pieces of drink bottles. On the other hand, the optimal cultivation pH was 7, while the optimal temperature was 23 °C, consistent with the average temperature of the sample collection area.

*A. guillouiae* I-MWF showed minimal growth when pure terephthalic acid (TPA), the aromatic monomers of PET, was used as the sole carbon source. Indeed, at neutral pH, TPA is a non-permeable molecule, and its diffusion through the cell membrane is unlikely, as computational studies have supported (Vermaas et al., 2019). In addition, we could not identify any putative TPA transporter gene in the *A. guillouiae* I-MWF genome, although genes encoding for benzoate/H(+) symporter (*benE*) and mayor facilitator superfamily (MFS) transporter are present (Table 5). However, it is important to highlight that little is known about TPA transporters in general. Despite that, *A. guillouiae* I-MWF has putative genes involved in the protocatechuate acid pathway, which can be linked to the metabolism of phenolic compounds (Table 5). Hypothetically, perhaps other PET depolymerization products, such as MHET and BHET, which are significantly more hydrophobic than TPA, could cross the cell membrane through diffusion.

The depolymerization product profile of the extracellular enzymes of *A. guillouiae* I-MWF is like other bacterial PETases and cutinases. The main product is MHET. For example, in *Ideonella skaieneis*, *Is*PETase produces mainly MHET, and a second extracellular enzyme, MHETase, is necessary to convert MEHT to TPA. Then, a specialized transporter in the periplasm imports the TPA into the cytoplasm (Yoshida et al., 2016). MHET is also the main product when marine PET2 (Wagner-Egea et al., 2023) and leaf-branch compost cutinase (LCC) from bacterial origin are used for TPA depolymerization (Tournier et al., 2020). The extracellular enzymatic extract from *Stenotrophomonas maltophilia* also yields MEHT as the major PET powder degradation product (Din et al., 2023). The fungal cutinase HiC (HiCut) and *Rhizomucor miehei* lipase (RmL) produced TPA as the dominant PET degradation product (Aristizábal-Lanza et al., 2022; Brackmann et al., 2023), while *Candida antarctica lipase* (CaL) produced mainly BHET (Brackmann et al., 2023). However, the product’s ratio profile could vary depending on factors such as incubation time, temperature and enzyme/polymer ratio, among others (Wagner-Egea et al., 2021).

The FT-IR analysis of the remaining PET, exposed to *A. guillouiae* I-MWF growth, has unveiled progressive chemical alterations over time, indicative of biodegradation. Notably, the peaks corresponding to C=C (1644.35 cm^-1^) and O-H have become increasingly pronounced as time has advanced. Specifically, the peak at 1644.35 cm^-1^, representing ν=(C=C), has gradually enhanced. As compared to untreated abiotic controls, the FT-IR spectra of PET film subjected to microbial treatment displayed the emergence of new peaks. After seven days of treatment with the *A. guillouiae* I-MWF strain, samples revealed the presence of two distinctive peaks associated with O-H and C=C functional groups. The heightened intensity of the O-H peak suggests the breakdown of the polymer. These observed changes in the presence or absence of functional groups are closely linked to microbial activities, providing crucial insights into the mechanism of plastic biodegradation (Auta et al., 2017).

In genome of *A. guillouiae* I-MWF identified the presence of at least 18 genes whose products are the extracellular hydrolases (Table 3). Genome is lacking cutinase-like enzymes but rich in putative lipases and esterases. Indeed, these groups of enzymes catalyses the ester bond hydrolysis, but they have different substrate specificities and functions. Cutinases can hydrolyse PET, and some of them are also known as PETases. Esterases, lipases, cutinases and PETases contain a serine residue as nucleophile, which is crucial in the catalysis, also histidine that acts as a base, and glutamate or aspartate that acts as proton donor. Thus, this catalytic triad works in a coordinated manner during the enzymatic reaction.

The molecular models obtained correspond to extracellular lipases and esterases predicted from the genome analysis of *A. guillouiae* I-MWF. All share the typical α/β-fold domain of hydrolases. They are diverse regarding the molecular weights 20 to 47 kDa and pÍs suggesting different functions. The catalytic triad is well-conserved in all models. Compared with cutinases, the modelled enzymes have CAP and lid domains in the case of lipase and a sharper catalytic site in the esterases. These structural features suggest that their catalytic interaction with PET can be affected by esteric hindrances. However, several lipases have been reported to be active against PET. For example, the highly promiscuous lipase B from *Candida antartica* (CALB), which natural substrates are triacyl glycerides, can also hydrolases polyesters such as PET oligomers (Świderek et al., 2023). Lipases from *Pseudomonas chlororaphis* have shown hydrolytic activity on petroleum-based polymers, such as poly-(ε-caprolactone) (PCL) and polyethene succinate (PES) (Mohanan et al., 2022).

A comparative study has shown that a lipase from *Thermomyces lanuginosus* produced slightly higher amounts of larger fragments from PET than the cutinase cutinases from *Thermobifida fusca* and *Fusarium solani* (Verma et al., 2021). *Acinetobacter* is a genus characterized by a wide variety of genes encoding lipases. Indeed, *A. guillouiae* I-MWF genome encodes a variety of extracellular lipases and esterases that could depolymerize partially powder PET. Future studies on the individual gene cloning and characterization of these enzymes would clarify their potential role in PET degradation.

## Conclusions

A bacterial strain capable of degrading PET powders was successfully isolated from the activated sludge. Following analysis of the 16S rDNA sequence and employing phylogenetic methods, it was identified as *Acinetobacter guillouiae*.

This study revealed the partial powder PET degradation of *A. guillouiae* I-MWF in a minimal salt medium. This was confirmed by FT-IR analysis, which detected alterations in the functional groups of PET when exposed to the bacteria. Moreover, HPLC-UV and LC-MS analyses identified depolymerization products, substantiating the biodegradation process and the strain’s capacity to utilize PET.

Although the bacterial strain has been successfully isolated, the enzymes responsible for PET degradation are still being identified and characterized. The sequenced genome comprises 4,178 coding genes, including 18 proteins with potential roles in PET biodegradation, spanning several putative lipases and three esterases. Future research, such as proteomics and gene cloning, aimed at elucidating the roles of enzymes in PET depolymerization, may lead to a deeper understanding of the underlying molecular mechanisms driving the observed biodegradation of PET powder.

## Acknowledgements

We thank Dr Ping Wang for supplying the powder PET. The Physiographic Society in Lund and The Nilsson-Ehle Foundation, Sweden, founded this project. Lund University provided open-access publication funding. The Higher Education Commission of Pakistan (HEC) has provided funds for the internships of Naheed Akhtar at Lund University, Sweden, under the International Research Support Initiative Program (IRSIP).

## Author contributions statement

Conceptualization: Javier A. Linares-Pastén, Naheed Akhtar, Afef Najjari

Formal analysis: Naheed Akhtar, Javier A. Linares-Pastén, Afef Najjari, Cecilia Tullberg, Carl Grey, Baozhong Zhang.

Funding acquisition: Javier A. Linares-Pastén.

Investigation: Naheed Akhtar, Afef Najjari, Javier A. Linares-Pastén.

Methodology: Naheed Akhtar, Afef Najjari, Cecilia Tullberg, Javier A. Linares-Pastén.

Project administration: Javier A. Linares-Pastén

Resources: Javier A. Linares-Pastén, Baozhong Zhang, Naheed Akhtar.

Supervision: Javier A. Linares-Pastén, Zahid Majeed, Muhammad Siddique Awan

Validation: Naheed Akhtar, Afef Najjari, Cecilia Tullberg, Javier A. Linares-Pastén, Carl Grey, Baozhong Zhang.

Writing – original draft: Naheed Akhtar, Afef Najjari, Javier A. Linares-Pastén,

Writing – review & editing: Javier A. Linares-Pastén, Afef Najjari, Naheed Akhtar, Zahid Majeed, Baozhong Zhang, Carl Grey.

## References

1. Aristizábal-Lanza, L., Mankar, S.V., Tullberg, C., Zhang, B., Linares-Pastén, J.A., 2022. Comparison of the enzymatic depolymerization of polyethylene terephthalate and AkestraTM using Humicola insolens cutinase. Frontiers in Chemical Engineering 4.

2. Auta, H., Emenike, C., Fauziah, S., 2017. Screening of Bacillus strains isolated from mangrove ecosystems in Peninsular Malaysia for microplastic degradation. Environmental Pollution 231, 1552–1559.

3. Bankevich, A., Nurk, S., Antipov, D., Gurevich, A.A., Dvorkin, M., Kulikov, A.S., Lesin, V.M., Nikolenko, S.I., Pham, S., Prjibelski, A.D., 2012. SPAdes: a new genome assembly algorithm and its applications to single-cell sequencing. Journal of computational biology 19, 455–477.

4. Bolger, A.M., Lohse, M., Usadel, B., 2014. Trimmomatic: a flexible trimmer for Illumina sequence data. Bioinformatics 30, 2114–2120.

5. Brackmann, R., de Oliveira Veloso, C., de Castro, A.M., Langone, M.A.P., 2023. Enzymatic post-consumer poly (ethylene terephthalate)(PET) depolymerization using commercial enzymes. 3 Biotech 13, 135.

6. Buchfink, B., Xie, C., Huson, D.H., 2015. Fast and sensitive protein alignment using DIAMOND. Nature methods 12, 59–60.

7. Buchholz, P.C., Feuerriegel, G., Zhang, H., Perez_Garcia, P., Nover, L.L., Chow, J., Streit, W.R., Pleiss, J., 2022. Plastics degradation by hydrolytic enzymes: The plastics_active enzymes database—PAZy. Proteins: Structure, Function, and Bioinformatics 90, 1443–1456.

8. Danso, D., Schmeisser, C., Chow, J., Zimmermann, W., Wei, R., Leggewie, C., Li, X., Hazen, T., Streit, W.R., 2018. New insights into the function and global distribution of polyethylene terephthalate (PET)-degrading bacteria and enzymes in marine and terrestrial metagenomes. Applied and environmental microbiology 84, e02773–02717.

9. Din, S.U., Satti, S.M., Uddin, S., Mankar, S.V., Ceylan, E., Hasan, F., Khan, S., Badshah, M., Beldüz, A.O., Çanakçi, S., Zhang, B., Linares-Pastén, J.A., Shah, A.A., 2023. The Purification and Characterization of a Cutinase-like Enzyme with Activity on Polyethylene Terephthalate (PET) from a Newly Isolated Bacterium Stenotrophomonas maltophilia PRS8 at a Mesophilic Temperature. Applied Sciences 13, 3686.

10. Donelli, I., Freddi, G., Nierstrasz, V.A., Taddei, P., 2010. Surface structure and properties of poly-(ethylene terephthalate) hydrolyzed by alkali and cutinase. Polymer degradation and stability 95, 1542–1550.

11. Farris, J.S., 1972. Estimating phylogenetic trees from distance matrices. The American Naturalist 106, 645–668.

12. Felsenstein, J., 1985. Confidence limits on phylogenies: an approach using the bootstrap. evolution 39, 783–791.

13. Gambarini, V., Pantos, O., Kingsbury, J.M., Weaver, L., Handley, K.M., Lear, G., 2022. PlasticDB: a database of microorganisms and proteins linked to plastic biodegradation. Database 2022.

14. Gasteiger, E., Hoogland, C., Gattiker, A., Duvaud, S.e., Wilkins, M.R., Appel, R.D., Bairoch, A., 2005. Protein identification and analysis tools on the ExPASy server. Springer.

15. Grant, J.R., Stothard, P., 2008. The CGView Server: a comparative genomics tool for circular genomes. Nucleic acids research 36, W181–W184.

16. Hiraga, K., Taniguchi, I., Yoshida, S., Kimura, Y., Oda, K., 2019. Biodegradation of waste PET: A sustainable solution for dealing with plastic pollution. EMBO reports 20, e49365.

17. Kang, Y.-S., Jung, J., Jeon, C.O., Park, W., 2011. Acinetobacter oleivorans sp. nov. is capable of adhering to and growing on diesel-oil. The Journal of Microbiology 49, 29–34.

18. Kelley, L.A., Mezulis, S., Yates, C.M., Wass, M.N., Sternberg, M.J., 2015. The Phyre2 web portal for protein modeling, prediction and analysis. Nature protocols 10, 845–858.

19. Kim, H.R., Lee, C., Shin, H., Kim, J., Jeong, M., Choi, D., 2023. Isolation of a polyethylene-degrading bacterium, Acinetobacter guillouiae, using a novel screening method based on a redox indicator. Heliyon 9.

20. Kumar, S., Stecher, G., Li, M., Knyaz, C., Tamura, K., 2018. MEGA X: molecular evolutionary genetics analysis across computing platforms. Molecular biology and evolution 35, 1547.

21. Kwon, G.-S., Kim, D.-G., Lee, J.-B., Shin, K.-S., Kum, E.-J., Sohn, H.-Y., 2006. Isolation of Acinetobacter calcoaceticus BP-2 capable of degradation of bisphenol A. Journal of Life Science 16, 1158–1163.

22. Lagesen, K., Hallin, P., Rødland, E.A., Stærfeldt, H.-H., Rognes, T., Ussery, D.W., 2007. RNAmmer: consistent and rapid annotation of ribosomal RNA genes. Nucleic acids research 35, 3100–3108.

23. Lai, J., Huang, H., Lin, M., Xu, Y., Li, X., Sun, B., 2023. Enzyme catalyzes ester bond synthesis and hydrolysis: The key step for sustainable usage of plastics. Front. Microbiol 14, 1113705.

24. Lefort, V., Desper, R., Gascuel, O., 2015. FastME 2.0: a comprehensive, accurate, and fast distance-based phylogeny inference program. Molecular biology and evolution 32, 2798–2800.

25. Li, C., Liu, C., Li, R., Liu, Y., Xie, J., Li, B., 2022. Biodegradation of dibutyl phthalate by the new strain Acinetobacter baumannii DP-2. Toxics 10, 532.

26. Liu, Z., Xie, W., Li, D., Peng, Y., Li, Z., Liu, S., 2016. Biodegradation of phenol by bacteria strain Acinetobacter calcoaceticus PA isolated from phenolic wastewater. International journal of environmental research and public health 13, 300.

27. Lowe, T.M., Eddy, S.R., 1997. tRNAscan-SE: a program for improved detection of transfer RNA genes in genomic sequence. Nucleic acids research 25, 955–964.

28. Lux, M., Krüger, J., Rinke, C., Maus, I., Schlüter, A., Woyke, T., Sczyrba, A., Hammer, B., 2016. acdc– Automated Contamination Detection and Confidence estimation for single-cell genome data. BMC bioinformatics 17, 1–11.

29. Marten, E., Müller, R.-J., Deckwer, W.-D., 2005. Studies on the enzymatic hydrolysis of polyesters. II. Aliphatic–aromatic copolyesters. Polymer degradation and stability 88, 371–381.

30. Meier-Kolthoff, J.P., Göker, M., 2019. TYGS is an automated high-throughput platform for state-of-the-art genome-based taxonomy. Nature communications 10, 2182.

31. Mohanan, N., Wong, C.H., Budisa, N., Levin, D.B., 2022. Characterization of polymer degrading lipases, LIP1 and LIP2 from Pseudomonas chlororaphis PA23. Frontiers in Bioengineering and Biotechnology 10, 854298.

32. Nemec, A., Musílek, M., Šedo, O., De Baere, T., Maixnerova, M., van der Reijden, T.J., Zdráhal, Z., Vaneechoutte, M., Dijkshoorn, L., 2010. Acinetobacter bereziniae sp. nov. and Acinetobacter guillouiae sp. nov., to accommodate Acinetobacter genomic species 10 and 11, respectively. International Journal of Systematic and Evolutionary Microbiology 60, 896-903.

33. Parte, A.C., Sardà Carbasse, J., Meier-Kolthoff, J.P., Reimer, L.C., Göker, M., 2020. List of Prokaryotic names with Standing in Nomenclature (LPSN) moves to the DSMZ. International Journal of Systematic and Evolutionary Microbiology 70, 5607–5612.

34. Perez-Garcia, P., Chow, J., Costanzi, E., Gurschke, M., Dittrich, J., Dierkes, R.F., Molitor, R., Applegate, V., Feuerriegel, G., Tete, P., 2023. An archaeal lid-containing feruloyl esterase degrades polyethylene terephthalate. Communications Chemistry 6, 193.

35. Pettersen, E.F., Goddard, T.D., Huang, C.C., Couch, G.S., Greenblatt, D.M., Meng, E.C., Ferrin, T.E., 2004. UCSF Chimera—a visualization system for exploratory research and analysis. Journal of computational chemistry 25, 1605–1612.

36. Richter, M., Rosselló-Móra, R., Oliver Glöckner, F., Peplies, J., 2016. JSpeciesWS: a web server for prokaryotic species circumscription based on pairwise genome comparison. Bioinformatics 32, 929–931.

37. Rodriguez-R, L.M., Konstantinidis, K.T., 2014. Bypassing cultivation to identify bacterial species. Microbe 9, 111–118.

38. Saitou, N., Nei, M., 1987. The neighbor-joining method: a new method for reconstructing phylogenetic trees. Molecular biology and evolution 4, 406–425.

39. Salvador, M., Abdulmutalib, U., Gonzalez, J., Kim, J., Smith, A.A., Faulon, J.-L., Wei, R., Zimmermann, W., Jimenez, J.I., 2019. Microbial genes for a circular and sustainable bio-PET economy. Genes 10, 373.

40. Świderek, K., Velasco-Lozano, S., Galmés, M.À., Olazabal, I., Sardon, H., López-Gallego, F., Moliner, V., 2023. Mechanistic studies of a lipase unveil effect of pH on hydrolysis products of small PET modules. Nature communications 14, 1–10.

41. Taniguchi, I., Yoshida, S., Hiraga, K., Miyamoto, K., Kimura, Y., Oda, K., 2019. Biodegradation of PET: current status and application aspects. Acs Catalysis 9, 4089–4105.

42. Tatusov, R.L., Galperin, M.Y., Natale, D.A., Koonin, E.V., 2000. The COG database: a tool for genome-scale analysis of protein functions and evolution. Nucleic acids research 28, 33–36.

43. Tatusova, T., DiCuccio, M., Badretdin, A., Chetvernin, V., Nawrocki, E.P., Zaslavsky, L., Lomsadze, A., Pruitt, K.D., Borodovsky, M., Ostell, J., 2016. NCBI prokaryotic genome annotation pipeline. Nucleic acids research 44, 6614–6624.

44. Teufel, F., Almagro Armenteros, J.J., Johansen, A.R., Gíslason, M.H., Pihl, S.I., Tsirigos, K.D., Winther, O., Brunak, S., von Heijne, G., Nielsen, H., 2022. SignalP 6.0 predicts all five types of signal peptides using protein language models. Nature biotechnology 40, 1023–1025.

45. Throne-Holst, M., Wentzel, A., Ellingsen, T.E., Kotlar, H.-K., Zotchev, S.B., 2007. Identification of novel genes involved in long-chain n-alkane degradation by Acinetobacter sp. strain DSM 17874. Applied and environmental microbiology 73, 3327–3332.

46. Torena, P., Alvarez_Cuenca, M., Reza, M., 2021. Biodegradation of polyethylene terephthalate microplastics by bacterial communities from activated sludge. The Canadian Journal of Chemical Engineering 99, S69–S82.

47. Tournier, V., Topham, C., Gilles, A., David, B., Folgoas, C., Moya-Leclair, E., Kamionka, E., Desrousseaux, M.-L., Texier, H., Gavalda, S., 2020. An engineered PET depolymerase to break down and recycle plastic bottles. Nature 580, 216–219.

48. Van Dexter, S., Boopathy, R., 2019. Biodegradation of phenol by Acinetobacter tandoii isolated from the gut of the termite. Environmental Science and Pollution Research 26, 34067–34072.

49. Verma, S., Meghwanshi, G.K., Kumar, R., 2021. Current perspectives for microbial lipases from extremophiles and metagenomics. Biochimie 182, 23–36.

50. Vermaas, J.V., Dixon, R.A., Chen, F., Mansfield, S.D., Boerjan, W., Ralph, J., Crowley, M.F., Beckham, G.T., 2019. Passive membrane transport of lignin-related compounds. Proceedings of the National Academy of Sciences 116, 23117–23123.

51. Wagner-Egea, P., Aristizábal-Lanza, L., Tullberg, C., Wang, P., Bernfur, K., Grey, C., Zhang, B., Linares- Pastén, J.A., 2023. Marine PET Hydrolase (PET2): Assessment of Terephthalate-and Indole-Based Polyester Depolymerization. Catalysts 13, 1234.

52. Wagner-Egea, P., Tosi, V., Wang, P., Grey, C., Zhang, B., Linares-Pastén, J.A., 2021. Assessment of Is PETase- Assisted Depolymerization of Terephthalate Aromatic Polyesters and the Effect of the Thioredoxin Fusion Domain. Applied Sciences 11, 8315.

53. Wang, P., Linares-Pastén, J.A., Zhang, B., 2020a. Synthesis, molecular docking simulation, and enzymatic degradation of AB-type indole-based polyesters with improved thermal properties. Biomacromolecules 21, 1078–1090.

54. Wang, Y., Coleman-Derr, D., Chen, G., Gu, Y.Q., 2015. OrthoVenn: a web server for genome wide comparison and annotation of orthologous clusters across multiple species. Nucleic acids research 43, W78–W84.

55. Wang, Z., Xin, X., Shi, X., Zhang, Y., 2020b. A polystyrene-degrading Acinetobacter bacterium isolated from the larvae of Tribolium castaneum. Science of the Total Environment 726, 138564.

56. Wei, R., Zimmermann, W., 2017. Microbial enzymes for the recycling of recalcitrant petroleum_based plastics: how far are we? Microbial biotechnology 10, 1308–1322.

57. Wingett, S.W., Andrews, S., 2018. FastQ Screen: A tool for multi-genome mapping and quality control. F1000Research 7.

58. Xu, L., Dong, Z., Fang, L., Luo, Y., Wei, Z., Guo, H., Zhang, G., Gu, Y.Q., Coleman-Derr, D., Xia, Q., 2019. OrthoVenn2: a web server for whole-genome comparison and annotation of orthologous clusters across multiple species. Nucleic acids research 47, W52–W58.

59. Yin, C.-F., Xu, Y., Zhou, N.-Y., 2020. Biodegradation of polyethylene mulching films by a co-culture of Acinetobacter sp. strain NyZ450 and Bacillus sp. strain NyZ451 isolated from Tenebrio molitor larvae. International Biodeterioration & Biodegradation 155, 105089.

60. Yoshida, S., Hiraga, K., Takehana, T., Taniguchi, I., Yamaji, H., Maeda, Y., Toyohara, K., Miyamoto, K., Kimura, Y., Oda, K., 2016. A bacterium that degrades and assimilates poly (ethylene terephthalate). Science 351, 1196–1199.

61. Yue, F., Zhang, L., Wang, J., Zhou, Y., Ye, B., 2017. Biodegradation of phthalate esters by a newly isolated Acinetobacter sp. strain LMB-5 and characteristics of its esterase. Pedosphere 27, 606–615.

62. Zhang, H., Lu, Y., Wu, H., Liu, Q., Sun, W., 2023. Effect of an Acinetobacter pittobacter on low-density polyethylene. Environmental Science and Pollution Research 30, 10495–10504.

63. Zrimec, J., Kokina, M., Jonasson, S., Zorrilla, F., Zelezniak, A., 2021. Plastic-degrading potential across the global microbiome correlates with recent pollution trends. MBio 12, e02155–02121.

